# Spatial multi-omics unveils the monoclonal origin, neuroendocrine plasticity, and microenvironment niches in combined small cell lung cancer

**DOI:** 10.64898/2026.01.31.702982

**Authors:** Zhuo Wang, Qing Luo, Jie Wu, Li Lu, Wenjie Ding, Yichun Zhao, Yongfeng Yu, Ruoran Qiu, Ling Zhu, Xinxing Ouyang, Wendi Xuzhang, Shun Lu, Wei Wei, Qihui Shi, Ziming Li

## Abstract

Combined small-cell lung cancer (cSCLC) is a rare and aggressive subtype of small-cell lung cancer (SCLC) characterized by mixed histology comprising SCLC and non-small cell lung cancer (NSCLC) or large cell neuroendocrine carcinoma (LCNEC) components. Despite its histological heterogeneity and even poorer prognosis than de novo SCLC, cSCLC is clinically managed as pure SCLC, largely due to the lack of molecular insights into its biology, lineage plasticity, and tumor microenvironment (TME). Here, we perform multi-omics profiling, including spatially-resolved whole-exome sequencing (WES), spatial transcriptomics (ST) and single-nucleus RNA sequencing (snRNA-seq), across 19 treatment-naïve cSCLC tumors spanning all major histological subtypes. Our analysis reveals that SCLC and NSCLC/LCNEC components share a monoclonal origin, with histological divergence characterized by distinct mutation and copy number alteration patterns. ST and snRNA-seq uncover spatially exclusive or interspersed tumor domains, with distinct TME compositions and immune landscapes. Notably, fibroblast-rich regions enriched for an aggressive fibroblast subtype form boundaries between tumor domains, potentially influencing immune TME and treatment responses. We identify extensive lineage plasticity within cSCLC, including active LUAD-to-SCLC transdifferentiation and SCLC subtype coexistence, suggesting transitional cellular states not captured by traditional diagnostics. Leveraging these insights, we developed the cSCLC Detector, a sensitive mutation-based diagnostic assay that improves the detection of cSCLC in tissue and liquid biopsy samples. Our findings offer critical insights into cSCLC lineage plasticity, cellular evolution, and microenvironmental interactions, underscoring the need for tailored treatment strategies and diagnostic frameworks for this aggressive cancer subtype.

## Introduction

Combined small-cell lung cancer (cSCLC) is a rare and poorly understood subtype of small cell lung cancer (SCLC), first identified by the World Health Organization in 1981,^1^ and represents approximately 2-5% of all SCLC cases in the literature over the past two decades.^2,3^ cSCLC is defined by a pathological mixture of SCLC and any subtype of non-small cell lung cancer (NSCLC) components, including lung adenocarcinoma (LUAD) and lung squamous cell carcinoma (LUSC), or large cell neuroendocrine carcinoma (LCNEC) subtype.^1^ The diagnosis of cSCLC is primarily established on pathological evaluation of surgically resected tumor specimens. cSCLC is often underdiagnosed in advanced stage unresectable patients because small biopsies may not capture the full histological diversity of the tumor.^2-6^

Traditionally, the mixed histologies within cSCLC were thought to arise from independent tumor populations, but recent genomic studies have challenged this notion, revealing that different histological components share common genetic mutations.^3,7-10^ This evidence suggests that cSCLC originates from a single progenitor cell that undergoes divergent differentiation, rather than representing two clonally distinct malignancies. Despite containing a mixture of SCLC and NSCLC (or LCNEC) components, cSCLC is treated as pure SCLC without specialized treatment protocol in current clinical practice.^2-6^ Clinical trials and treatment guidelines do not typically distinguish between pure SCLC and cSCLC due to its rarity and the lack of large, subtype-specific pathological and molecular studies. Notably, cSCLC is associated with an even poorer prognosis than *de novo* SCLC,^11,12^ yet the mechanisms driving its aggressive nature remain elusive.

One emerging concept in cSCLC research is that the coexistence of different histological components reflects underlying cancer cell plasticity rather than being a simple admixture of two distinct carcinoma types. This concept has been particularly well illustrated in cSCLC cases with SCLC/LUAD mixed histology, where a recent study by Quintanal-Villalonga et al.^13^ demonstrated that the mixed histology reflects LUAD-to-SCLC transformation driven by predisposed molecular alterations, such as *TP53*/*RB1* function loss, that prime LUAD tumor cells for neuroendocrine (NE) differentiation.^10^ By analyzing LUAD/SCLC cases with matched pre-transformation LUAD, post-transformation SCLC, never-transformed LUAD, and *de novo* SCLC, that study provided strong evidence that molecular determinants of NE transformation are already present before histological conversion occurs in treatment-naïve cSCLC samples with SCLC/LUAD histology, regardless of *EGFR* mutation. They also performed comprehensive multi-omics profiling of combined SCLC/LUAD tumors, establishing that the distinct histological components share a monoclonal origin and identifying key transcriptional reprogramming events that drive NE transformation. However, while these findings advance our understanding of cellular plasticity in cSCLC, existing studies are scarce and largely phenotypic or based on bulk-level sequencing, which have significant limitations. For example, bulk sequencing lacks the resolution to determine whether NE differentiation occurs as a gradual shift in tumor composition, where NSCLC cells are progressively replaced by SCLC-like cells, or whether individual tumor cells transition through intermediate/hybrid states before fully acquiring an SCLC phenotype. Furthermore, bulk profiling cannot resolve the spatial organization of transformed tumor components or reveal how different components within cSCLC interact with immune and stromal cells within the tumor microenvironment (TME), a critical factor in understanding tumor evolution and treatment response.

Understanding the clonal origins, lineage and molecular plasticity, and TME organization of cSCLC is critical for refining diagnostic criteria and developing effective therapeutic strategies tailored to its unique biology. To this end, we utilize spatially resolved genomic and transcriptomic sequencing, coupled with single-cell RNA sequencing,^14-19^ to 19 treatment-naïve cSCLC patient samples spanning all three major histological subtypes, including SCLC/LUAD, SCLC/LUSC, and SCLC/LCNEC. By characterizing the transcriptional states, clonal evolution, and spatial heterogeneity of cSCLC, we provide insights into how lineage plasticity is regulated in cSCLC and uncover TME factors that may influence molecular phenotype, immune landscape, and treatment responses.

## Results

### cSCLC patient cohort and study design

A total of 19 treatment-naïve cSCLC patients were enrolled in this study, including 10 SCLC/LUAD (P1-P10), 3 SCLC/LUSC (P11-P13) and 6 SCLC/LCNEC (P14-P19) patients (Figure 1A, Table S1). To date, this is the cSCLC cohort with the highest proportion of SCLC/LUAD patients (>50%).^2-9^ In each tumor sample, the areas of SCLC and NSCLC (or LCNEC) components were clearly defined by pathological examinations (Figure 1B and Figure S1), except for P17, who had SCLC and LCNEC components intimately mixed without clear boundary. In the formalin-fixed and paraffin-embedded (FFPE) cSCLC samples, multiple regions of SCLC and NSCLC (or LCNEC) components were collected by microdissection for multi-region whole exome sequencing (WES). In total, WES was performed on 95 SCLC, NSCLC and LCNEC regions from 16 patients, with an average of 6 regions sequenced per patient (Figure 1C). Single-nucleus RNA sequencing (snRNA-seq) and spatial transcriptomics (ST) analyses were conducted on 12 and 6 patients, respectively, encompassing all three histological subtypes.

**Figure 1.**
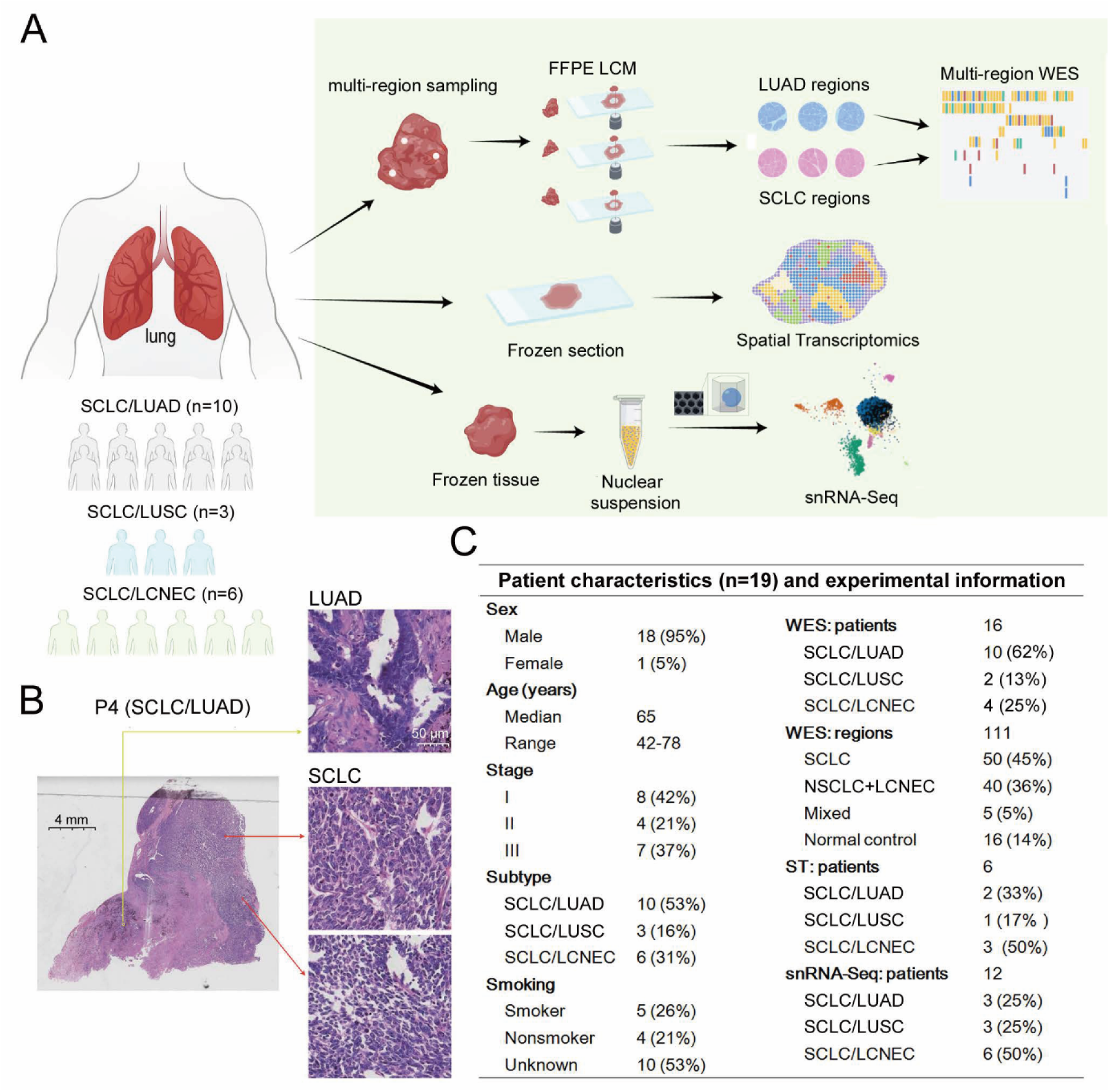
Overview of study design, methods and patient cohort. (A) Schematic illustration of experimental workflow and multi-omic characterization of cSCLCs. The study involved multi-region WES, spatial transcriptomics (ST) and snRNA-seq. (B) Representative H&E staining showing the coexistence of SCLC and LUAD histological components within the same tissue sample from P4. (C) Demographics, clinical characteristics, and summary of multi-omics data generation for the 19 cSCLC patients enrolled in this study.

### Mutational landscape of cSCLC via multi-region WES

Multi-region WES was used to spatially characterize mutation profiles of SCLC and NSCLC (or LCNEC) components within the same FFPE sections of cSCLC tumors with the paired normal adjacent tissue. A total of 95 tumor regions (50 SCLC, 40 NSCLC+LCNEC and 5 mixed regions) and 16 paired normal adjacent tissues were collected by microdissection from 16 patients and sequenced at high depth (Table S2), ensuring robust data quality and coverage. The average number of somatic mutations was 547 (range: 85-1133) for tumor regions (Table S3).

The most recurrently mutated genes across patients were *TP53* and *RB1* (Figures 2A and S2). *TP53* somatic variants were present in all SCLC and NSCLC (or LCNEC) regions of all the cSCLC patients except for P3, with all the regions from each patient sharing the identical variants (Table S4), establishing them as clonal mutations of cSCLC. *RB1* mutations were identified as clonal mutations in 6 patients and as subclonal mutations in P2 and P9. In P2, *RB1* was found to be mutant in 5 regions (4 SCLC, 1 LUAD) but wild-type in only 1 LUAD region (Figures 2A). *EGFR* mutations, which are frequent driver mutations in LUAD, were detected in 40% (4/10) of cSCLC patients with SCLC/LUAD subtype. In these four patients, *EGFR* mutations were detected in all SCLC and LUAD regions of P6, P8, and P9, establishing them as clonal mutations of cSCLC. In P7, *EGFR* mutations were present in 3 out of 4 of SCLC regions and all 2 LUAD regions. The types of *EGFR* mutations detected in cSCLC patients with the SCLC/LUAD subtype were similar to those reported in pure LUAD patients, including in-frame deletion of exon 19 (19Del) in P6 and P9, L858R point mutation in P7, and a combination of E709V and G719C in P8 (Figure 2B, Table S5). Importantly, the 40% frequency of *EGFR* mutations, shared across both SCLC and LUAD components of this subtype, was found consistent with those reported from two independent Asian SCLC/LUAD cohorts (Figure 2B, Table S5).^11,14^ This frequency is close to the 44% *EGFR* mutation frequency reported for Asian male patients with pure LUAD, but significantly higher than that observed in pure *de novo* SCLC patients (∼3%),^10^ suggesting unique genetic characteristics of the SCLC components in cSCLC and supporting the NE transformation within the SCLC/LUAD subtypes.

**Figure 2.**
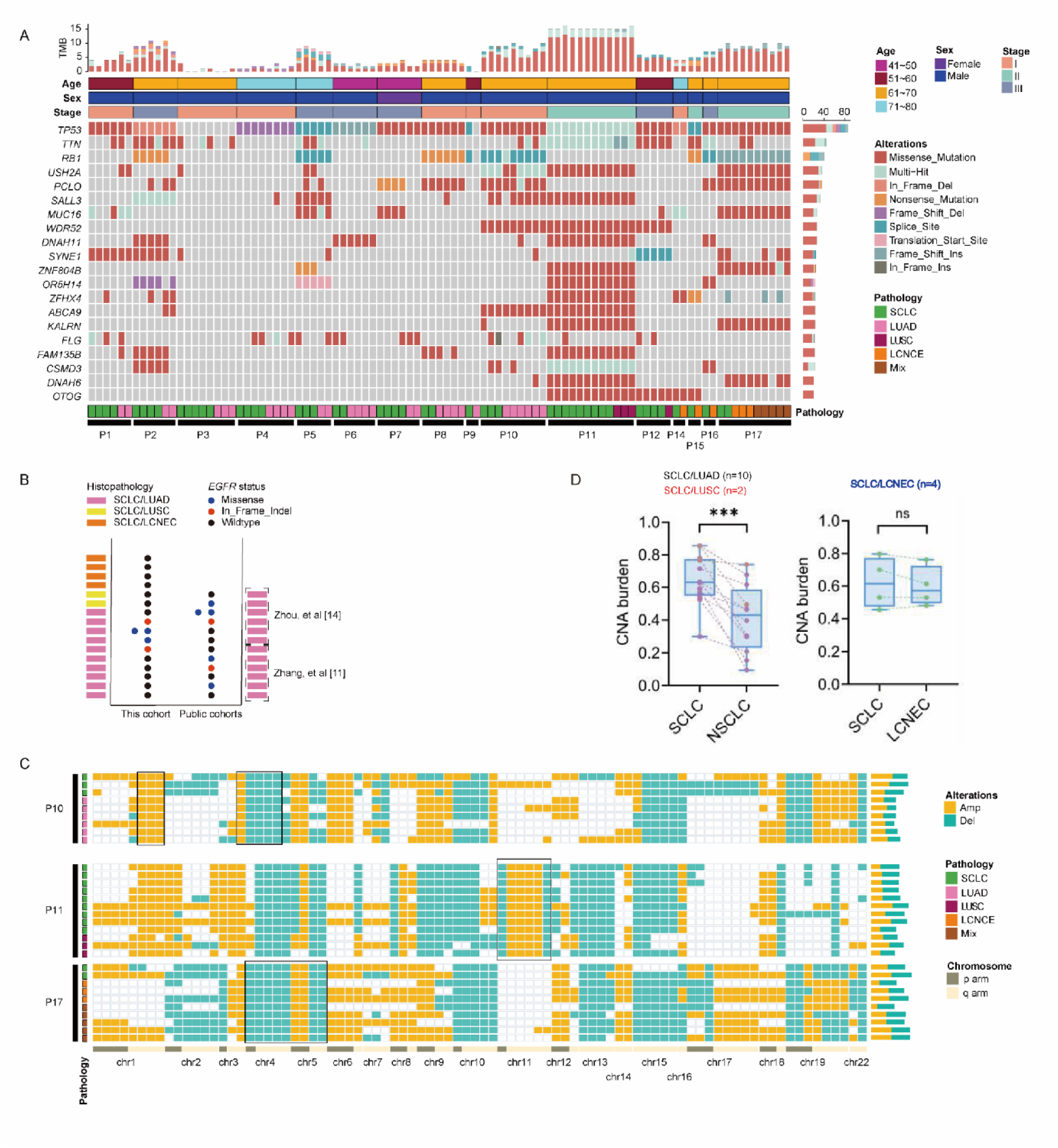
Mutational landscape and copy number alterations of cSCLC via multi-region WES. (A) Oncoplot displaying the somatic mutation landscape of cSCLC patients. The heatmap shows the top 20% most frequently mutated genes in different regions from cSCLC patients. Clinical characteristics (age, sex, stage) and tumor mutational burden (TMB) are annotated at the top. The bar plots summarize the mutation counts per gene (right) and per sample (top). (B) Prevalence of *EGFR* mutations in cSCLC samples. The dot plot compares *EGFR* mutation status (missense, in-frame indel, or wildtype) across the cohort in this study and two additional public cSCLC cohorts.^11,14^ Colors indicate the histopathological subtype (SCLC/LUAD, SCLC/LUSC or SCLC/LCNEC). See also Table S5. (C) Heatmap showing arm-level CNA across multiple tumor regions from 3 representative cSCLC patients (P10, P11, and P17). Rows represent individual tumor regions, and columns represent chromosome arms (1p–22q). CNA are color-coded as amplifications (yellow) and deletions (teal); gray indicates no call. The pathology track (left) denotes the histological subtype of each sequenced region. The right bar plot shows the CNA burden for each region, expressed as the fraction of chromosome arms altered, stratified by amplifications (yellow) and deletions (teal). Black frames highlight consistent altered chromosome arms across regions in individual patients. (D) Comparison of averaged CNA burdens between SCLC and NSCLC (combined LUAD and LUSC, n=12) or LCNEC (n=4) components within cSCLC patients. Data are presented as boxplots where the central line represents the median, box limits indicate the interquartile range (25th to 75th percentiles), and whiskers extend to the minimum and maximum values. Pairwise comparisons were performed using a two-sided Wilcoxon rank-sum test. \*\*\**p* < 0.001; ns, not significant.

For genome-wide copy number alteration (CNA), different histological subtypes from the same patients showed largely consistent CNA patterns, implicating that they may be derived from the same progenitor cancer cells (Figure 2C). The most frequently gained chromosome arms in cSCLC were 1q, 5p, and 6p, while losses were commonly detected in the 3p, 4q, 5q, 10q and 15q chromosome arms (Figures 2C and S3). The frequent CNA signatures were consistent in the SCLC and NSCLC (or LCNEC) components of cSCLC, and were different from pure SCLCs that were usually amplified at 3q and 5q and losses at 13q (containing *RB1*) and 17p (containing *TP53*).^11,14^ Notably, the loss of 3p has been identified as a risk factor for NE transformation in SCLC/LUAD subtype.^13^ Consistently, we observed 3p loss in 6 out of 10 SCLC/LUAD samples, in 1 out of 2 SCLC/LUSC samples, and in all 4 SCLC/LCNEC samples sequenced. Interestingly, CNA burdens in the SCLC regions were significantly higher than those in the LUAD or LUSC regions (Figure 2D). However, no significant difference of CNA burdens was found in SCLC and LCNEC regions of SCLC/LCNEC samples (Figure 2D), possibly due to the higher proliferation and higher genomic instability associated with NE histology.

### Genomic evolution reveals monoclonal origin of cSCLC

We investigated mutational profiles of multiple SCLC and NSCLC (or LCNEC) regions from the same cSCLC tumors and found high concordance of somatic mutations across different histological regions of interest (ROIs), even those isolated from different tissue blocks. For example, we collected three tumor blocks (B1-B3) from P11 and sequenced 2 SCLC regions and 1 LUSC region from block 1 (B1), 3 SCLC regions and 1 LUSC region from block 2 (B2), 4 SCLC regions and 1 LUSC region from block 3 (B3) with WES. Venn diagrams show large overlaps of somatic mutations between ROIs from different histological subtypes within the same block and across different blocks (Figure 3A), indicating high concordance of somatic mutations from SCLC and LUSC regions from different locations of the tumor, suggesting a monoclonal origin of these different histological subtypes. Similar concordance of somatic mutations was also identified in SCLC and LUAD regions across different blocks from P8 (Figure S4).

**Figure 3.**
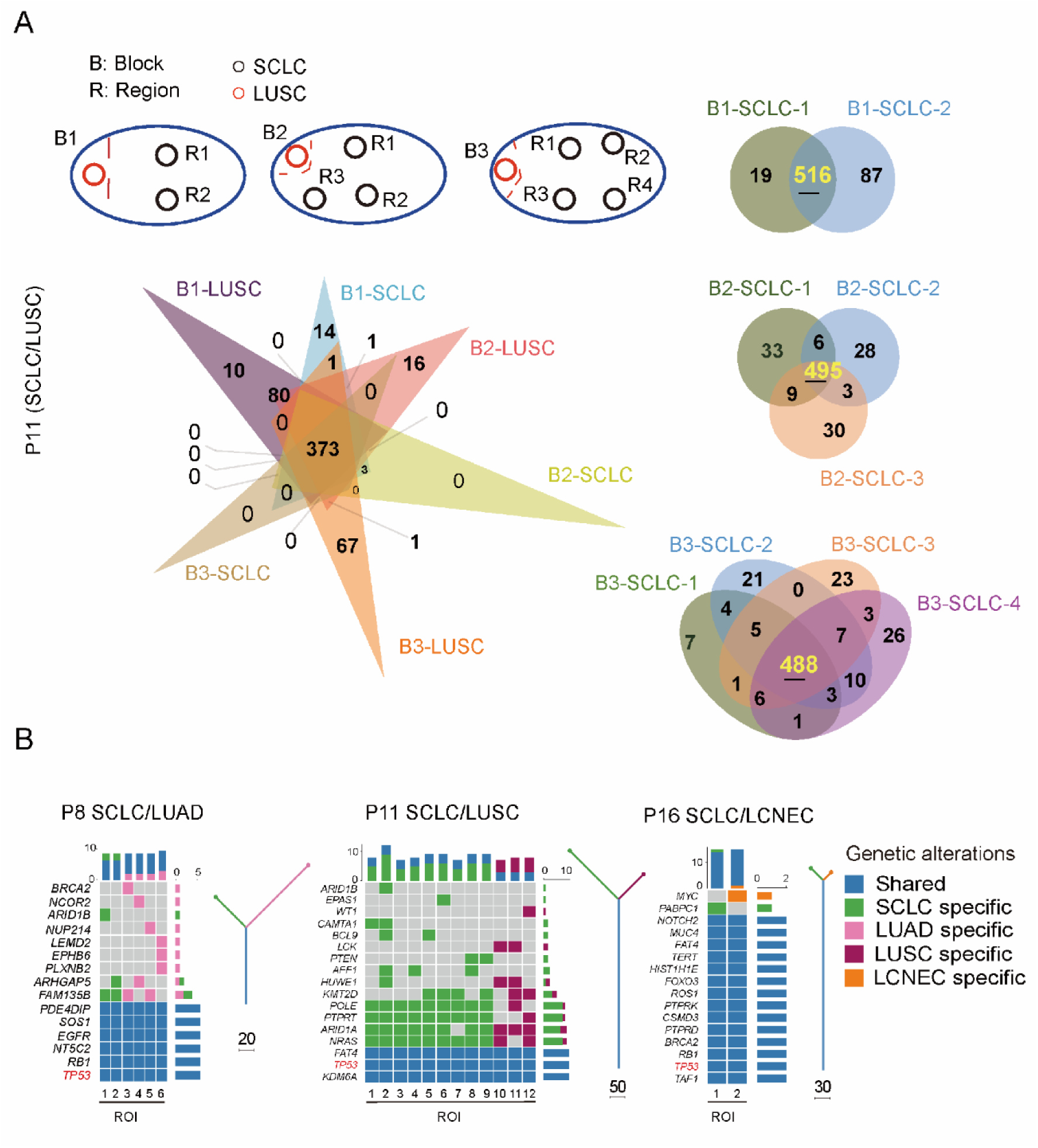
Clonal evolution and phylogenetic analysis of cSCLC via multi-region WES. (A) Consistency of somatic mutations across different spatial regions in P11. Top left: Schematic illustration of tissue blocks (B1–B3) and spatial regions (R) sampled via laser capture microdissection (LCM). Bottom left: Venn diagram displaying the overlap of somatic mutations across different tissue blocks and histological components, highlighting a core set of 373 shared mutations. Right: Venn diagrams illustrating the consistency of somatic mutations among different SCLC regions within specific tissue blocks (B1, B2, and B3). (B) Phylogenetic trees and mutational heatmaps of three representative cSCLC patients: P8 (SCLC/LUAD), P11 (SCLC/LUSC), and P16 (SCLC/LCNEC). For each patient, the central heatmap displays nonsynonymous somatic mutations in selected cancer genes across regions of interest (ROIs). Mutations are color-coded by regional specificity: shared/truncal (blue), SCLC-specific (green), LUAD-specific (pink), LUSC-specific (purple), and LCNEC-specific (orange). The top stacked bar plots indicate the total mutation burden per ROI, partitioned by the same specificity categories. Phylogenetic trees (right) depict the inferred evolutionary relationships and monoclonal origin of the components. The trunk represents early clonal mutations shared across all ROIs (with the number of truncal mutations indicated), while branches represent lineage-specific diversification.

To elucidate the clonal evolution of cSCLC, we constructed phylogenetic trees using nonsynonymous mutations identified in different histological ROIs of each sample (Table S3). Our analysis revealed that multiple histological tumor ROIs within each sample shared many oncogenic truncal mutations, such as *TP53, RB1, EGFR*, *PTEN,* suggesting that they occurred before the histologic divergence (Figures 3B and S5). In the intensively sampled SCLC/LUSC case P11, truncal alterations in *TP53* and the chromatin regulators *KDM6A*^20^ and *FAT4* were detected in both SCLC and LUSC components, whereas early driver events such as *NRAS* and *ARID1A,*^21^ a key epigenetic tumor suppressor gene in lung cancer, showed near-ubiquitous distribution across regions (*NRAS* in all SCLC ROIs and 2 of 3 LUSC ROIs; *ARID1A* in all LUSC ROIs and 8 of 9 SCLC ROIs), with the few mutation-negative ROIs likely reflecting lower tumor cellularity and variant allele fractions below the detection threshold associated with spatial heterogeneity in tumor content. By contrast, mutations in *AFF1*, *PTEN*, *BCL9*, and *CAMTA1* were confined to SCLC ROIs, while *LCK* and *WT1* mutations were restricted to LUSC ROIs, illustrating lineage-specific private branches that emerged after divergence from the common progenitor. Similar patterns of shared truncal driver alterations together with lineage-restricted private mutations were observed throughout the cohort (Figures 3B and S5). Notably, the key driver mutation *TP53* was consistently found at the root of the phylogenetic trees across nearly all the samples, in line with the almost universal *TP53* alterations found in both SCLC and NSCLC regions of the cSCLC samples from previous reports.^14^ This recurrent combination of shared truncal driver alterations and lineage-restricted private mutations is best explained by a branched evolutionary model in which morphologically distinct SCLC and NSCLC (or LCNEC) components arise from a single ancestral clone that diverged into distinct morphologic and lineage states over time. Importantly, in four SCLC/LUAD patients (P6–P9), we detected *EGFR* mutations—common in LUAD but extremely rare in *de novo* SCLC—in both SCLC and LUAD regions (Figure 2B and Table S5). This finding suggests that SCLC and LUAD cells originated from a common *EGFR*-mutant progenitor, further supporting the role of LUAD-to-SCLC NE transformation in cSCLC with mixed SCLC/LUAD histology.

### Mutation-based diagnostic test for cSCLC detection with improved sensitivity

The diagnosis of cSCLC is primarily based on pathological evaluation of surgically resected tumor specimens. In most cSCLC tumors, the SCLC and NSCLC (or LCNEC) components are spatially separated, with the SCLC regions often being substantially larger. Given the limited sampling inherent to biopsy procedures, there is a risk that the NSCLC component may not be captured, leading to the misclassification of cSCLC as *de novo* SCLC in unresectable patients who rely on biopsy-based diagnosis. To this end, we developed a mutation-based diagnostic test (cSCLC Detector) for sensitive detection of cSCLC from SCLC patients using tissue biopsy or liquid biopsy such as blood. The cSCLC Detector is a 4-gene SNV panel (*EGFR*, *KRAS*, *BRAF*, *PIK3CA*). All genetic alterations selected for the panel were reported to be highly specific to LUAD and LUSC, and have been identified as truncal mutations shared by SCLC components either in our samples or in previous cSCLC studies (Figure 4A, Tables S6 and S7).^22^ To assess the performance of this 4-gene panel detector, we collected a total of 38 cSCLC samples (SCLC/LUAD and SCLC/LUSC) that were confirmed by histopathology, including 12 samples from this study and 26 samples from three cohorts in previous studies.^8,11,14^ cSCLC Detector showed 61% sensitivity in this cohort (n=38), and particularly, 74% and 27% sensitivity in SCLC/LUAD (n=27) and SCLC/LUSC (n=11) patients, respectively (Figure 4B, Table S6). The relatively lower sensitivity for the SCLC/LUSC subtype is primarily attributable to the lack of highly specific and recurrent driver mutations in LUSC. We then tested the cSCLC Detector with tissue-based sequencing data of two previously considered *de novo* SCLC cohorts (n=64), and identified cSCLC-Detector-positive results in 14.1% of samples (9/64, Figure 4C, 4D, Table S6).^11,14^ Additionally, cSCLC Detector was further tested with blood circulating tumor DNA (ctDNA) data from another cohort of previously considered de novo SCLC patients (n=254) and showed cSCLC-Detector–positive results in 14.2% of samples (36/254), similar to the fraction from tissue-based genomic sequencing (Figure 4C, 4D, Tables S6 and S7).^23^ Compared with previous studies reporting approximately 2-5% cSCLC cases in all SCLC patients,^2,3^ cSCLC Detector enables molecular detection of cSCLC through small-sized biopsies with improved sensitivity, which could be attributed to the detection of these NSCLC-specific genes even when only SCLC components are sampled (Figure 4E, Tables S6 and S7). Additionally, blood ctDNA-based cSCLC Detector may provide a non-invasive method for molecular cSCLC detection.

**Figure 4.**
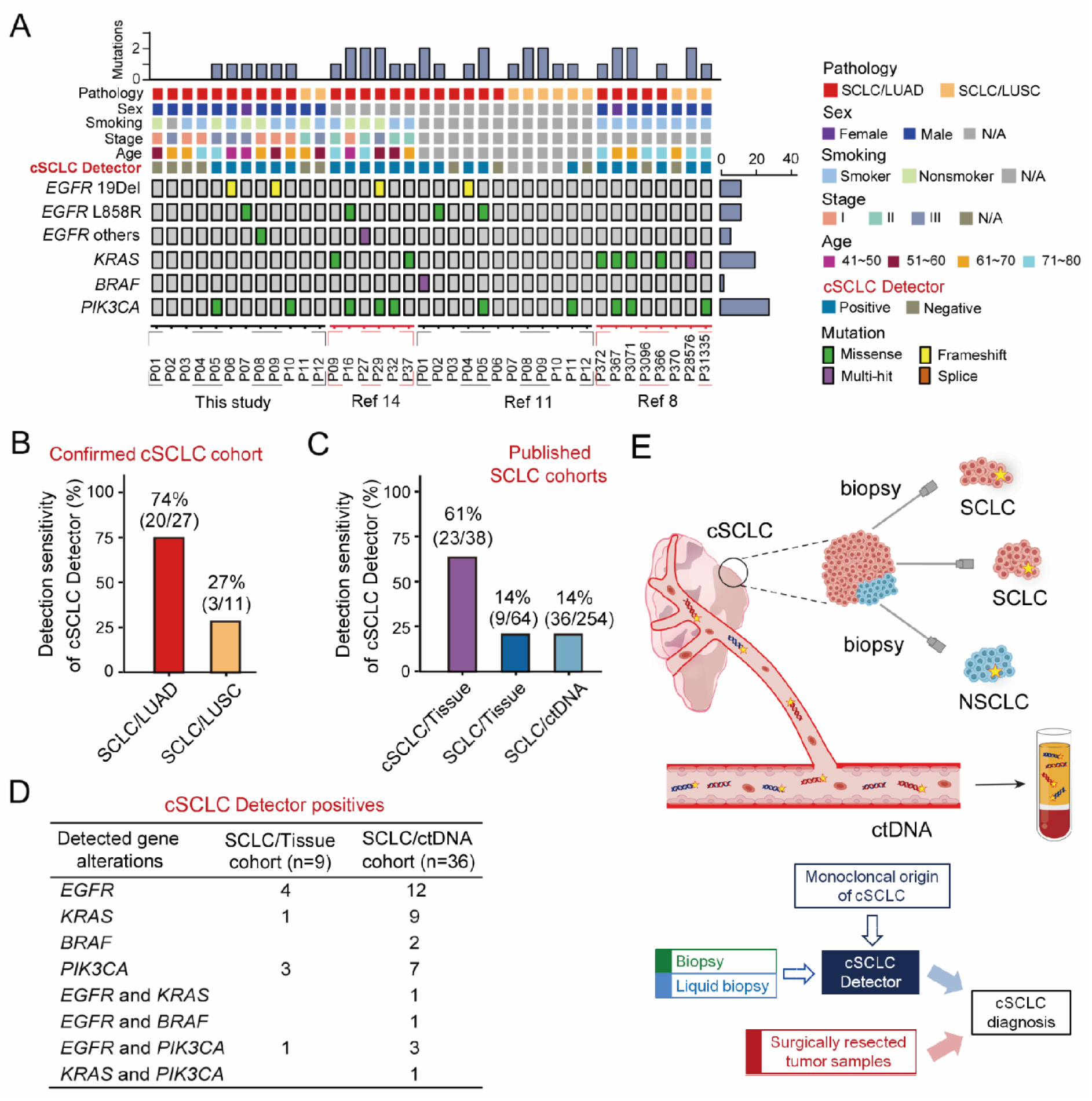
Development and validation of the cSCLC detection for enhanced diagnostic sensitivity. (A) Landscape of somatic mutations detected by the cSCLC Detector in a combined cohort of 38 cSCLC patients. The cohort comprises patients from this study (n=12) and three external public datasets (n=26).^8,11,14^ See also Tables S7. The oncoplot displays the mutation status of four NSCLC-driver genes (*EGFR*, *KRAS*, *BRAF*, and *PIK3CA*) alongside clinical characteristics and cSCLC Detector results (Positive/Negative). Top bar plots represent the mutation count per patient, and right bar plots show the mutation frequency per gene across the cohort. (B) Performance of the cSCLC Detector for cSCLC detection in the SCLC/LUAD (n=27), SCLC/LUSC (n=11) subgroups and all tissue samples of confirmed cSCLC cohort (n=38). (C) Performance of the cSCLC Detector for cSCLC detection in the published SCLC cohort (n=64), as well as blood samples of SCLC cohort (n=254). (D) Summary of gene mutations detected with cSCLC Detector panel in the "positive" cases from the published SCLC tissue (n=9) and ctDNA (n=36) cohorts shown in (C). (E) Schematic illustration addressing the limitations of small biopsy-based pathological evaluation for cSCLC diagnosis and the principle of cSCLC Detector for cSCLC detection with improved sensitivity.

### Spatial transcriptomics identifies tumor domains segregated by aggressive cancer-associated fibroblasts

To further investigate molecular features of different histological subtypes and their interactions within the TME of cSCLC, we performed spatial transcriptomics (ST) on tissue sections from 6 patients with spatial spots ranging from 1,186 to 2,556 per sample. We integrated ST spots from all six samples into a single Uniform Manifold Approximation and Projection (UMAP) embedding, identifying ten distinct clusters (Figure 5A). To annotate these clusters, we curated a reference scRNA-seq dataset comprising published data from tumor samples of patients with LUAD, LUSC, and major SCLC subtypes (SCLC-A: *ASCL1*-dominant; SCLC-N: *NEUROD1*-dominant; SCLC-P: *POU2F3*-dominant).^24,25^ This unified reference dataset was employed consistently throughout our analysis for both cluster annotation and subsequent ST deconvolution. Spot annotations were assigned by evaluating similarity to reference annotations using a combined similarity metric that integrated Jensen-Shannon divergence and cosine similarity (Figure 5B).

**Figure 5.**
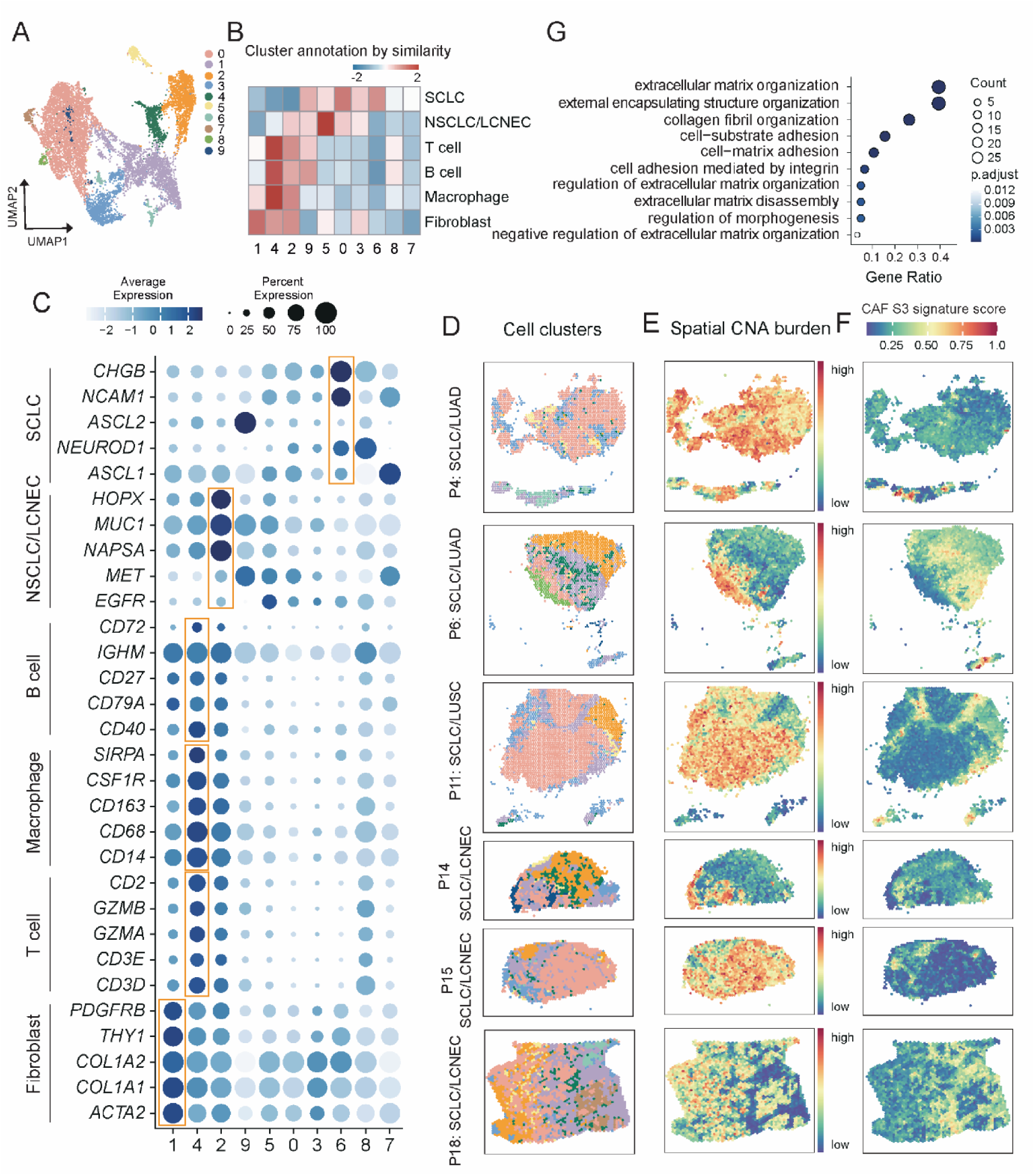
Co-embedding of spatial transcriptomics spots for characterizing cSCLC microenvironment. (A) UMAP embedding of integrated spatial transcriptomics (ST) spots from 6 cSCLC samples, identifying 10 distinct transcriptional clusters based on global gene expression profiles. (B) Heatmap illustrating the cluster annotation strategy. Cluster identities were assigned by calculating the similarity (z-score) between ST spots and a reference scRNA-seq dataset (comprising SCLC, LUAD, and LUSC) using a combined metric of Jensen-Shannon divergence and cosine similarity.^24,25^ (C) Dot plot displaying the expression of canonical marker genes used to define cell lineages. The dot color encodes the average expression level of the gene within the cluster (see color scale). Dark blue indicates higher average expression, while light colors indicate lower expression. The dot size representss the percentage of cells expressing the gene within the cluster (see size scale). Larger dots correspond to a higher proportion of cells expressing the gene. (D) Spatial images depicting the location of identified clusters for 6 cSCLC patients. (E) Spatial CNA burden was quantified across different histological regions. The CNA burden is color coded. Warmer colors (red) indicate higher CNA burden. (F) The enrichment of the aggressive CAF S3 subtype signature in the fibroblast regions that spatially interpose and segment the tumor domains. High CAF S3 signature scores (red) spatially correspond to fibroblast regions that interpose and segment the tumor domains. (G) Gene Ontology (GO) enrichment analysis of genes defining the CAF S3 signature. Dot size indicates the number of genes enriched in the pathway, and color represents statistical significance (adjusted p-value).

In the clustering analysis, Cluster 1 (21.5% of the spots) was characterized by high expressions of *THY1*, *PDGFRB*, and collagen-related genes, indicating fibroblast-dominated regions (Figure 5B, 5C). Cluster 4 (5.9%) consisted of immune cells expressing marker genes such as *CD3D* (T cells), *CD79A* (B cells), and *CD68* (macrophages), indicating immune-infiltrated regions. Clusters 2 (13.4%) exhibit strong NSCLC/LCNEC signals with a mixture of fibroblasts and immune cells. In contrast, cluster 6 displays signatures associated with enrichment of SCLC cells, marked by *CHGB*, *NCAM1*, *NEUROD1* and *ASCL1* (Figure 5C). However, *POU2F3* expression was not detected in the SCLC enriched cluster, despite the presence of spatial regions characteristic of SCLC-P subtypes as shown below, potentially due to the inherent limitations of ST in capturing low-abundance transcripts. Other clusters displayed ambiguous identities, sharing molecular features with several tumor types, potentially derived from spatial spots where different cell types are highly intermixed.

We projected the defined clusters onto their spatial coordinates within the ST images, revealing distinct tumor and non-tumor domains across the tissue sections. In all cSCLC samples, separated tumor domains were displayed (Figure 5D). These spatial patterns aligned with CNA profiles, where tumor-enriched regions exhibited more pronounced genome-wide copy number changes (Figure 5E). Moreover, distinct tumor domains in cSCLC exhibited varying levels of immune infiltration. Cluster 2, which is enriched for NSCLC/LCNEC components, showed high immune infiltration based on immune marker expression (Figure 5B and 5C), whereas Cluster 6, enriched for SCLC components, displayed relatively lower immune infiltration, suggesting that the cellular composition of tumor domains may underlie spatial heterogeneity in immune exclusion within cSCLC.

Interestingly, the fibroblast domain (cluster 1, lavender spots) is frequently observed interposing between and segmenting distinct tumor domains within the cSCLC TME, highlighting a potential role for fibroblasts in spatially structuring the tumor landscape (Figure 5D). In cSCLC samples with clear SCLC/NSCLC (or LCNEC) compartmentalization, these boundary-localized fibroblasts coincide with collagen-rich stromal bands separating SCLC from NSCLC regions, resembling the COL11A1⁺ CAF barriers recently described in NSCLC.^26^A previous study reported that SCLC patients whose tumors are enriched for cancer-associated fibroblast subtype 3 (CAF S3), an aggressive subtype localized at the tumor edge of SCLC, exhibit significantly worse clinical outcomes.^27^ Our analysis showed that these tumor-domain-segmenting fibroblasts in cSCLC samples are enriched for CAF S3 subtype signature (Figures 5F and S6), marked by expression of extracellular matrix (ECM), collagen organization- and cell-matrix adhesion signatures (Figure 5G). Consistent with a COL11A1⁺ myCAF-like state, these fibroblasts express *COL11A1* together with multiple fibrillar collagens (*COL1A1*, *COL1A2*, *COL3A1*, *COL10A1*, *COL12A1*), ECM/activated-fibroblast markers (*FAP*, *POSTN*, *CTHRC1*, *GREM1*), and matrix-remodeling factors such as *THBS2*, *SPARC*, *MMP2*, and *MMP11*, indicating an ECM-producing, desmoplastic program that closely aligns with the previously defined COL11A1⁺ CAF gene module in NSCLC (Figure S6).

In contrast, fibroblasts residing within NSCLC-enriched domains (cluster 2) show an inflammatory CAF (iCAF)-like program characterized by *ADH1B*, *CXCL12*, *CXCL14*, *IL6*, and *PTGS2* expression and comparatively low levels of fibrillar collagens, mirroring the ADH1B⁺ iCAFs described in NSCLC (Figure S6).^26^ Notably, CAF S3 fibroblasts display high expression of pro-migratory genes such as *FNDC1*^28^ and *TEM4*^29^ (Figure S6), which have been implicated in immunosuppression and metastasis promotion. This spatial organization, in which a collagen-dense CAF S3/COL11A1⁺ band separates SCLC from adjacent NSCLC (or LCNEC) tumor domains and coincides with lymphocyte-poor SCLC regions, is consistent with a CAF-mediated, collagen-based immune exclusion mechanism analogous to that reported in NSCLC. The widespread enrichment of this aggressive CAF S3 subtype within the cSCLC TME across nearly all histological subtypes suggests that CAF S3 may contribute to the poorer prognosis of cSCLC, and may potentially drive its increased therapeutic resistance and facilitate disease progression compared to *de novo* SCLC. Although co-embedding analysis revealed interesting TME features within cSCLC, higher resolution analysis and additional validation are needed to quantitatively estimate the fractions of various cell types, as well as their spatial distribution and interactions.

### Spatial deconvolution reveals differential TME features of SCLC and NSCLC domains within cSCLC

To quantitatively estimate the cell type distribution within the TME, we proceeded with spatial deconvolution using the SPOTlight algorithm^30^ for each spot of the ST images. Spatial deconvolution utilized the same reference scRNA-seq dataset introduced earlier.^24,25^ To enhance the signal-to-noise ratio, we selected a group of marker genes that are differentially expressed for each cell type using the reference scRNA-seq data, establishing a reliable framework to distinguish distinct cell types (Figure S7A). To validate the robustness of this approach, we first tested it with a public 10X Visium ST dataset of LUAD and LUSC tissue sections.^31^ The deconvolution procedure accurately identified respective NSCLC tumor regions, fully consistent with the published pathological annotations and without confounding SCLC signals, confirming the high cell-type specificity of our deconvolution approach (Figure S7B).

Spatial deconvolution revealed spatial organization of tumor and non-tumor components, which closely matched the domains observed in the co-embedding analysis (Figures 6A, S8, S9 and S10). Tumor components were deconvolved into major SCLC subtypes (SCLC-A, -N, and -P) and NSCLC subtypes (LUAD and LUSC) or LCNEC subtype, while non-tumor components included fibroblasts, endothelial cells, and various immune cell types such as macrophages, T cells, and B cells. The SCLC and NSCLC (or LCNEC) components in the majority of samples (P6, P11, P14, P15) exhibited spatial exclusivity, forming well-defined, non-overlapping spatial regions (Figures 6A and S8), consistent with pathological observation, while displaying interspersed spatial distribution in remaining samples (P4, P18) (Figures S8-S10). This interspersed spatial organization might suggest a closer interaction or a partial NE transformation in the transcriptomic profiles between the two phenotypes.

**Figure 6.**
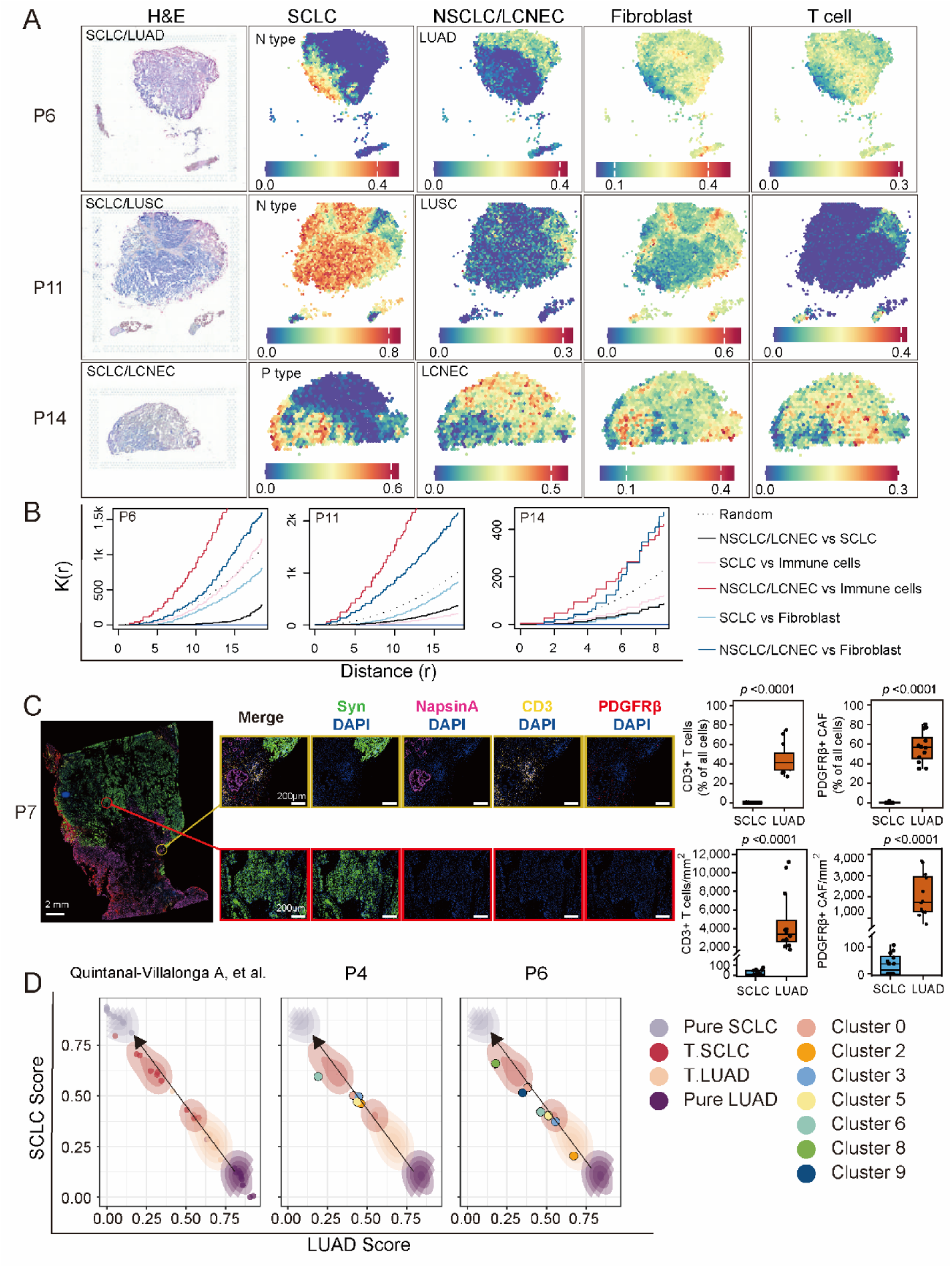
Spatial architecture and lineage plasticity suggesting *in situ* LUAD-to-SCLC transdifferentiation within cSCLS. (A) Spatial deconvolution of cell type compositions in 3 representative cSCLC cases: P6 (SCLC/LUAD), P11 (SCLC/LUSC), and P14 (SCLC/LCNEC). Left: H&E staining. Right: Spatial maps showing the inferred fractions of SCLC subtypes, NSCLC (or LCNEC) components (LUAD/LUSC/LCNEC), fibroblasts, and T cells from cell type deconvolution. (B) Ripley’s K-function plots showing spatial relationships between tumor components (SCLC and NSCLC) and infiltration components (immune and fibroblast cells). (C) Multiplex immunofluorescence (mIF) validation of the immune-excluded phenotype in SCLC regions. Left: Representative mIF images of patient P7 stained for Synaptophysin (SCLC), Napsin A (LUAD), CD3 (T cells), and PDGFRβ (CAFs). Yellow boxes indicate Napsin A^+^ LUAD domains, which are characterized by substantial infiltration of CD3^+^ T cells and PDGFRβ^+^ CAFs. Red boxes indicate Synaptophysin^+^ SCLC domains, which exhibit a "cold" immune phenotype with sparse T cell and fibroblast infiltration. Right: Quantification of CD3^+^ T cell and PDGFRβ^+^ CAF infiltration (percentage and density) in SCLC vs. LUAD regions (n=12 randomly selected ROIs). Box plots show the median (center line) and interquartile range (box); whiskers extend to 1.5×IQR. Statistical significance was determined by a two-sided Mann-Whitney U test. \**p*<0.0001; *p* values are indicated above the brackets. Scale bars: 2 mm (whole tissue, Left) and 200 μm (zoom, Right). (D) Projection of LUAD and SCLC transcriptional identity scores defined by quadratic programming. The left panel displays the reference dataset,^13^ showing the distribution of never-transformed LUAD (Pure LUAD, dark purple), pre-transformation LUAD (T.LUAD, light orange), post-transformation SCLC (T.SCLC, red), and *de novo* SCLC (Pure SCLC, light purple) on the plane defined by LUAD Score (x-axis) and SCLC Score (y-axis). The black arrow indicates the trajectory of neuroendocrine (NE) transformation. The middle and right panels show the projection of spatial transcriptomics spots from cSCLC patients P4 and P6, respectively, onto this transformation trajectory. Colored dots represent distinct spatial clusters.

Spatial deconvolution revealed remarkable differences between SCLC and NSCLC (or LCNEC) components in cSCLCs. Consistent with the co-embedding analysis, the deconvolution results showed that, for samples displaying spatial exclusivity, NSCLC/LCNEC regions were characterized by high infiltration of immune cells and fibroblasts, while SCLC regions exhibited minimal infiltration (Figures 6A, S8, S9 and S10). To quantify these spatial patterns, we applied Ripley’s K function^32^ to assess spatial relationships between tumor, immune, and stromal components (Figure 6B). The Ripley’s K function evaluates whether points exhibit aggregation, separation, or randomness. In samples where SCLC and NSCLC/LCNEC components were spatially exclusive, NSCLC/LCNEC regions exhibited stronger clustering with immune cells and fibroblasts, whereas SCLC regions demonstrated a tendency to repel infiltrating cells, indicating an immune-desert niche.

To experimentally validate the computationally inferred spatial segregation of tumor and TME components, we performed multiplex immunofluorescence (mIF) on tissue sections from seven cSCLC cases (P1-P7) of the SCLC/LUAD subtype. This subtype was selected because synaptophysin (Syn) and Napsin A allow for precise spatial discrimination between SCLC and LUAD lineages, respectively. We utilized lineage-specific markers to distinguish SCLC (Synaptophysin⁺) and LUAD (Napsin A⁺) components, alongside markers for T cells (CD3^+^) and cancer-associated fibroblasts (PDGFRβ^+^). Consistent with the spatial deconvolution and Ripley’s K analysis, mIF imaging revealed distinct, histology-dependent microenvironmental niches within single tumor specimens (Figure 6C). Napsin A⁺ LUAD domains were characterized by substantial infiltration of CD3⁺ T cells and PDGFRβ⁺ fibroblasts. In contrast, adjacent synaptophysin⁺ SCLC domains exhibited a "cold" immune phenotype with sparse T cell and fibroblast presence (Figures 6C, and S11). These orthogonal data confirm that the histological divergence in cSCLC is physically coupled with distinct immune and stromal landscapes, where the NSCLC component supports an infiltrated TME while the SCLC component maintains an immune-excluded niche, consistent with previous clinical findings.^33^

To investigate why these adjacent tumor compartments exhibit such divergent TMEs, we first asked whether the immune dichotomy could be explained by distinct oncogenic drivers. As shown above, multi-region WES revealed that SCLC and NSCLC (or LCNEC) components within each tumor share the same set of truncal driver alterations, arguing against genetically hard-wired differences in driver mutations as the primary cause of their contrasting immune states (Figures 3B and S5). We therefore turned to transcriptional and stromal features, focusing on three representative cases (P6, SCLC/LUAD; P11, SCLC/LUSC; and P14, SCLC/LCNEC), which together span the major cSCLC histologic subtypes and display clear spatial separation between SCLC and NSCLC (or LCNEC) domains. Voxel-level ST analysis in these tumors showed that SCLC-enriched regions consistently express lower levels of HLA class I genes (*HLA-A*, *HLA-B*, *HLA-C*) and *B2M* than adjacent NSCLC/LCNEC regions, suggesting impaired MHC-I antigen presentation as one contributor to T cell exclusion in the SCLC compartment (Figure S12). In parallel, fibroblast-rich bands at SCLC–NSCLC interfaces are strongly enriched for the CAF S3/COL11A1⁺ program and collagen-remodeling signatures (Figures 5 and S6), consistent with boundary-forming, desmoplastic CAFs capable of creating physical and biochemical barriers to immune infiltration.^26^ Together, these data indicate that the immune dichotomy in cSCLC may arise from a combination of lineage-specific downregulation of antigen presentation in SCLC regions and CAF S3–associated, collagen-dense stromal barriers at SCLC–NSCLC boundaries.

### Spatial transcriptomics reveals *in situ* LUAD-SCLC transdifferentiation

The histologic transformation of LUAD to an aggressive NE phenotype resembling SCLC was first observed in *EGFR*-mutant LUAD under selective therapeutic pressure.^34^ This phenomenon, a hallmark of lineage plasticity in lung cancer, was later found to occur independently of *EGFR* mutations, including in treatment-naïve settings.^14^ Given the monoclonal origin of the SCLC and NSCLC components in cSCLC, we sought to investigate how these components compare to pure SCLC and LUAD and where they reside within the NE transformation spectrum. To investigate the trajectory of neuroendocrine transformation specifically within the SCLC/LUAD subtype, we projected ST data from the SCLC and LUAD domains of our two SCLC/LUAD cases with spatial profiling (P4 and P6) onto a reference plane defined by LUAD and SCLC feature scores, derived from differentially expressed genes (DEGs) in our reference scRNA-seq dataset (see Method details).^24,25^

To contextualize these projections within the NE transformation trajectory, we incorporated transcriptomic data from a previous study that included never-transformed LUAD, *de novo* SCLC, histologic transformational LUAD and SCLC samples in the combined histology (T-LUAD and T-SCLC), and pretransformation LUAD samples that underwent NE transformation at a later stage and posttransformation SCLC samples that were previously diagnosed as LUAD.^13^ The resulting projection revealed a continuum along the diagonal axis, extending from never-transformed LUAD (lower right) to *de novo* SCLC (upper left), with T-LUAD and T-SCLC positioned in between, forming a clear trajectory of NE transformation (Figure 6D). Notably, ST data from SCLC and LUAD domains aligned more closely with T-LUAD and T-SCLC regions rather than the extremes of pure LUAD or SCLC, suggesting that these regions represent transitional plastic states with coexisting LUAD and SCLC transcriptomic signatures (Figure 6D). Interestingly, in P6, where SCLC and LUAD zones were spatially segregated, the projected transcriptome states of LUAD and SCLC spots were more polarized along the transformation spectrum. In contrast, in P4, where LUAD and SCLC components were spatially interspersed, the transcriptomic states of both domains were closer to the intermediate zone of the NE transformation trajectory, suggesting that a more spatially intermixed tumor architecture might be associated with an intermediate state with increased lineage plasticity actively shaping a transition between LUAD and SCLC. However, due to the large spot size in ST data, which captures multiple cells per spot, it remains unclear whether this mixed transcriptomic signature simply represents a gradual shift in tumor composition, with LUAD cells progressively replaced by SCLC-like cells, or whether individual tumor cells indeed transition through intermediate/hybrid states before fully adopting an SCLC phenotype. Resolving this question will require single-cell resolution analysis to pinpoint the precise mechanisms underlying NE plasticity in cSCLC.

### Single-nucleus RNA sequencing reveals distinct transcriptional states of cSCLC cells from pure SCLC and NSCLC

We performed snRNA-seq on 12 cSCLC samples (Figure 1C), and identified 150,738 high-quality cells in total. These data were integrated with the reference scRNA-seq dataset of pure SCLC and NSCLC samples mentioned above.^24,25^ The success of batch correction was evidenced by a correct overlap of immune cell and fibroblast components across different datasets (Figure 7A and 7B).^24,25^

**Figure 7.**
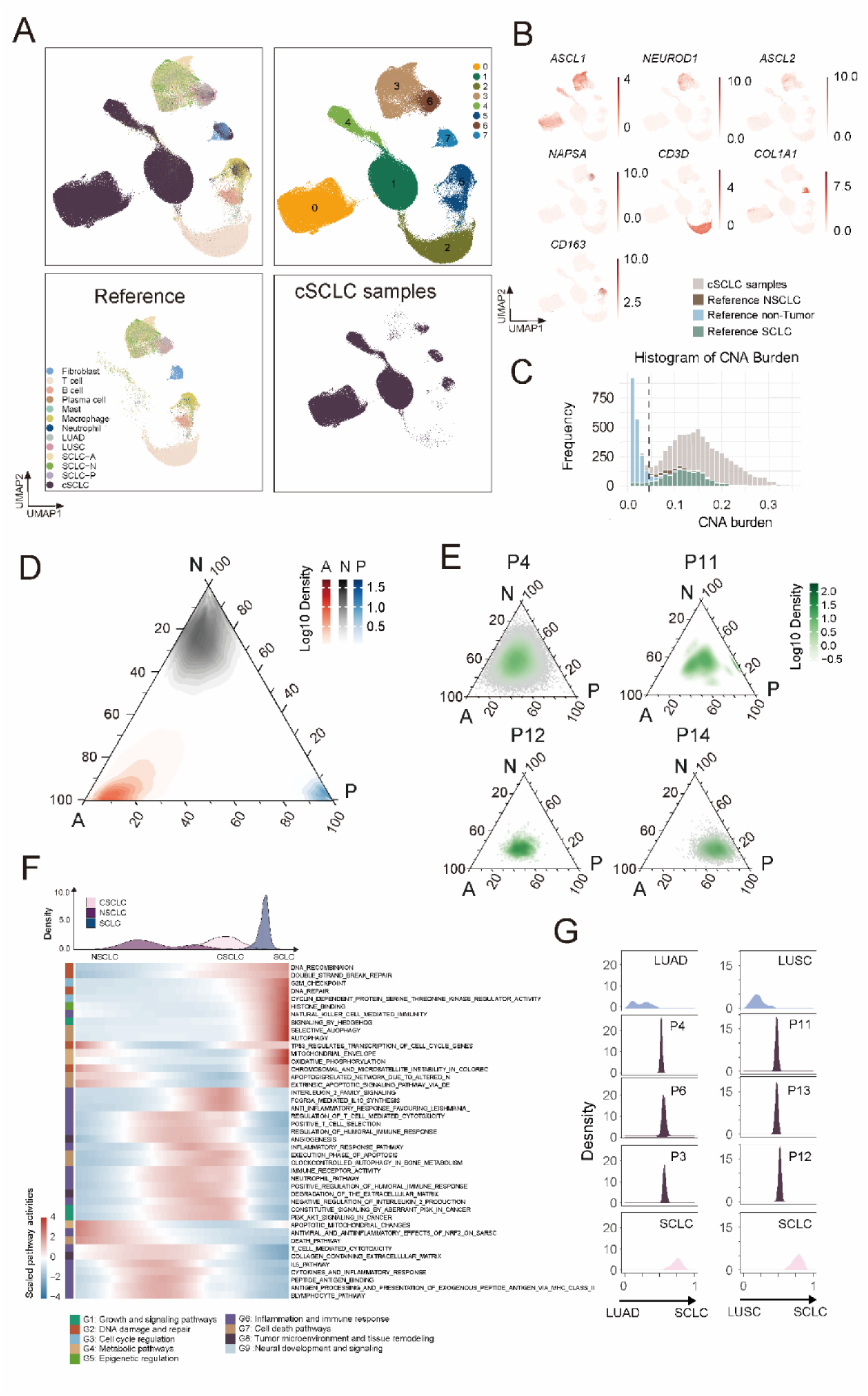
SnRNA-seq profiling identifies transcriptional plasticity of cSCLC tumor cells. (A) UMAP projection (upper left) of snRNA-seq data from cSCLC samples in this study and public datasets, and the cell clustering within the same UMAP (upper right). Separated embedding of the cells from the public reference datasets (lower left) and cSCLCs in this study (lower right) within the same UMAP. (B) Selected marker gene expressions on the UMAP from (A). (C) Histogram showing CNA burden distributions of tumor and non-tumor cells. (D) Ternary plots illustrating the distribution of pure SCLC cells across the A, N, and P subtype transcriptional space, showing distinct clustering toward their respective vertices (A in red, N in black, P in blue) based on normalized AUCell scores for subtype-specific gene sets. (E) Ternary plots of individual cSCLC samples (P4, P11, P12, P14), where each point represents a single SCLC-like cell and density contours (in green) highlight regions with high concentrations of cells. (F) Heatmap showing scaled pathway activity scores for individual cells ordered along a pseudotime-like trajectory spanning NSCLC, cSCLC, and SCLC states. Top: density plots showing the distribution of NSCLC, cSCLC, and SCLC cells along the NE transformation axis. Left: functional categories annotated by color. (G) Density plots showing the distribution of quadratic programming-derived SCLC-NSCLC identity scores for canonical LUAD, LUSC, and SCLC samples (top and bottom panels) as well as individual cSCLC patient samples (middle panels, labeled P3-P13). Each panel represents the distribution of single-cell identities of a specified patient projected onto a continuum from LUAD or LUSC (score = 0) to SCLC (score = 1), with intermediate values indicating a hybrid transcriptional state between NSCLC and SCLC.

Cell clustering identified distinct SCLC and NSCLC populations, with cluster 3 expressing SCLC markers such as *ASCL1*, *ASCL2*, and *NEUROD1*, and cluster 6 expressing NSCLC markers including *MUC1* and *NAPSA*. Fibroblasts formed cluster 7, characterized by *COL1A1* expression, while T cells grouped into cluster 2, marked by *CD3D* and *CD3E*. Cluster 5 comprised a mix of immune populations, including macrophages expressing *CD14* and *CD163* in its upper region, and B cells, plasma cells, neutrophils, and mast cells in its lower region, with B cell markers *CD79A* and *CD40* prominently expressed (Figures 7A, 7B, and S13).

Single nuclei isolated from cSCLC tumor samples showed substantial overlap with NSCLC, fibroblast, and macrophage clusters, but limited overlap with SCLC and T cell clusters. Additionally, the majority cells of cSCLC formed distinct clusters (clusters 0, 1, and 4), which were identified as malignant cells based on their significantly elevated genome-wide CNA burdens (Figure 7A and 7C). Examination of lung cancer markers showed that these cSCLC clusters expressed SCLC-associated genes such as *ASCL1*, *SCG2*, *ASCL2* (Figures 7B and S13), suggesting resemblance to pure SCLC cells but with distinct transcriptional states.

### High SCLC subtype plasticity and coexistence in cSCLC

SCLC molecular subtypes are characterized by mutually exclusive expression of *ASCL1*, *NEUROD1*, and *POU2F3*, representing A, N, and P types, respectively. Reanalysis of our reference scRNA-seq data^24,25^ from 21 pure *de novo* SCLC patients confirmed that *POU2F3* expression is essentially mutually exclusive with both *ASCL1* and *NEUROD1* (Figure S14A).^27,34^ SCLC-A cells express minimal *NEUROD1*, whereas a distinct subset of N-type cells exhibits low *ASCL1* levels (Figure S14A). These patterns suggest limited plasticity between A and N subtypes in pure SCLC, which is consistent with a recent report showing ∼10% of SCLC tumor cells co-expressing canonical markers from different molecular subtypes, primarily the NE lineage transcription factors, *ASCL1* and *NEUROD1*.^33^

In contrast, our ST deconvolution indicated co-localization of two subtypes in the majority of samples. For example, we observed co-localization of N-type and A-type SCLC cells in P6, P11, and P15, while P18 was dominated by A-type cells. (Figure S8). This is consistent with spots co-expressing high levels of *ASCL1* and *NEUROD1* (Figure S14B). In P15, *NEUROD1* expression was not detected by the spatial transcriptomics analysis. However, the SCLC-N transcriptional signature, defined by the differential marker genes of SCLC-N, was nevertheless evident and contributed to a substantial SCLC-N fraction in the spatial deconvolution (Figure S8). In P14, we observed coexistence of A- and P-type cells with higher P-type proportions (Figure S8); although *POU2F3* expression was low, likely due to transcript dropout in ST, other tuft cell-like P-type markers such as *ASCL2* and *MOCOS*^24^ (Figure S14B) were clearly expressed alongside *ASCL1*, indicating the presence of both A- and P-type subtypes in the TME of P14.

Our findings indicate that multiple subtype transcriptional signatures often coexist within the same spatial regions in cSCLC. However, it remains unclear whether this reflects a mixture of distinct subtype-specific cells within the same spots or the presence of cells in a hybrid plastic state, exhibiting molecular features of multiple pure subtypes. To resolve this, we assessed A, N, and P subtype plasticity in cSCLC tumor cells using the snRNA-seq data. We first identified SCLC-like cells by applying SPOTlight^30^ deconvolution to inferCNV-defined tumor cells (Figure 7C), classifying those with SCLC scores >0.75 and NSCLC scores <0.25 as SCLC-like single cells. We then quantified A, N, and P subtype activities in these cells by calculating enrichment scores for subtype-specific markers (Figure S7A), enabling us to represent each cell’s transcriptional identity within A-N-P subtype space. To establish a reference for comparison, we analyzed canonical single cells representing each SCLC subtype (A, N, P) from pure SCLC patients using the reference scRNA-seq dataset^24,25^ and visualized their distributions on a ternary plot (Figure 7D). These reference cells clustered tightly toward the respective A, N, or P vertices, confirming clear separation of pure subtypes.

In contrast, SCLC-like single cells from cSCLC samples (Figures 7E and S15) exhibited broad distributions across the ternary space, with many cells occupying intermediate compositional regions rather than clustering at a single subtype vertex. This pattern indicates substantial subtype plasticity in cSCLC, as individual SCLC-like tumor cells display hybrid expression signatures spanning multiple subtypes, a stark contrast to the mutually exclusive subtype profiles observed in pure SCLC tumor cells. Notably, this plasticity was not limited to neuroendocrine A and N subtypes; we also detected strong P-type signatures coexisting with A and/or N markers in individual cells, as exemplified by P14, which is consistent with ST deconvolution result (Figures 7E and S8). This suggests plasticity between NE and POU2F3 subtypes in SCLC-like cells of cSCLC, an observation not previously reported in pure SCLC samples.

To quantitatively estimate the fraction of cells in the subtype-hybrid states, we defined for each cell a dominance ratio as the maximum of its normalized A, N, and P subtype signature scores divided by their mean, such that high values indicate a single predominant subtype program and low values reflect co-activation of multiple programs. Using reference single-subtype SCLC cells from pure tumors to calibrate this metric, we set a hybrid cutoff at the mean dominance ratio minus two standard deviations, which classified only 3.7% of reference cells as hybrid, indicating that the threshold is conservative and rarely misclassifies *bona fide* single-subtype cells (Figure S16). Applying this threshold to cSCLC samples, 33% of SCLC-like tumor cells fell below the cutoff, indicating a markedly elevated fraction of transcriptionally hybrid cells in cSCLC compared with pure SCLC and quantitatively supporting the enhanced subtype plasticity inferred from the ternary representation. Moreover, these single-cell data reveal that genuine subtype plasticity within individual SCLC-like cells accounts for the presence of multiple subtype transcriptional signatures in the cSCLC spatial transcriptomics data, rather than a mere mixture of distinct subtype-specific cells within the same spatial spots.

We further confirmed these transcriptomic findings via immunohistochemistry (IHC) staining of the three major SCLC lineage transcription factors ASCL1, NEUROD1, and POU2F3 in selected cSCLC cases. In P6, ASCL1 and NEUROD1 proteins were co-localized within the same tumor regions, consistent with the A/N subtype co-localization inferred from ST and the co-expression of *ASCL1* and *NEUROD1* (Figures S14B, S17A and S17B). In P18, SCLC cells were uniformly ASCL1-positive and NEUROD1/POU2F3-negative, matching its A subtype-only ST profile (Figures S14B and S17A). In P15, by contrast, tumor cells showed strong ASCL1 positivity but lacked detectable NEUROD1 transcript or protein (Figures S14B and S17A), despite a clear SCLC-N transcriptional signature in the deconvolution analysis (Figure S8), illustrating that subtype programs can remain detectable at the gene-signature level even when individual marker genes fall below the sensitivity of ST or IHC. Finally, although matched ST data were not available for P19, its IHC profile-strong POU2F3 staining with absent ASCL1 and NEUROD1-mirrored the P-dominant position of its tumor cells in the A-N-P ternary single-cell map (Figures S15 and S17). Collectively, these protein-level patterns corroborate our ST and snRNA-seq subtype assignments and support the presence of both classical single-subtype regions and regions enriched for cells with overlapping subtype programs in cSCLC.

### cSCLC exhibit transcriptional intermediacy along the NSCLC-to-SCLC continuum

Given our observation of spots that display transcriptomic signatures intermediate between LUAD and SCLC in the SCLC/LUAD ST specimens (Figure 6D), we further leveraged this snRNA-seq data set to interrogate whether these cells occupy transcriptional states that are truly intermediate or merely mixed at the tissue level. Using SPOTlight^30^ deconvolution against the reference scRNA-seq dataset of pure NSCLC and SCLC samples,^24,25^ we computed cell-wise “NSCLC” and “SCLC” identity scores and ranked every cell along this continuum. As expected, cSCLC cells were broadly distributed between the poles of bona fide NSCLC and SCLC reference cells, confirming a unimodal spectrum of intermediate transcriptional identities (Figure 7F).

To interrogate how biological programs evolve along this axis, we curated eight functional pathway categories: growth and signaling pathways (G1), DNA damage and repair (G2), cell cycle regulation (G3), metabolic pathways (G4), epigenetic regulation (G5), inflammation and immune response (G6), cell death pathways (G7), and TME and tissue remodeling (G8) from MSigDB.^35,36^ Single-cell pathway activities were quantified using AUCell^37^ and analyzed across cells ordered by their relative NSCLC-SCLC identity scores, which revealed coordinated, monotonic shifts for several modules. For example, cells with high NSCLC scores displayed strong enrichment T cell–mediated cytotoxicity, antigen-presentation and neutrophil chemotaxis, in line with the highly inflamed microenvironment typical of NSCLC tumors.^38^ Concomitantly, extracellular-matrix (ECM) organization and collagen-remodeling signatures peaked in these cells, echoing matrisome-focused studies that emphasize stromal re-engineering as a hallmark of NSCLC progression. Moving rightwards along the continuum, we observed a gradual and ultimately sharp rise in G2 (DNA recombination, double-strand break repair) and G3 (G2/M checkpoint, cyclin-dependent kinase regulation) activities, culminating in classical SCLC cells, highlighting a shift toward more proliferative state with increased genomic instability. Multiple reports indicate that the ubiquitous loss of *TP53* and *RB1* and a dependency on DNA-damage-response (DDR) pathways in SCLC fuels hyper-proliferation yet renders the tumor vulnerable to PARP, ATR or CHK1 inhibition.^39,40^ The NSCLC-to-SCLC trajectory was marked by a switch from cytokine-driven inflammatory metabolism to heightened oxidative phosphorylation (OXPHOS) and selective autophagy (Figure 7F). Recent studies show that SCLC cells leverage mitochondrial respiration and fatty-acid β-oxidation for rapid growth, whereas autophagy confers chemoresistance via NRBF2-p62 interactions.^41,42^ Our data echo these findings: OXPHOS and autophagy scores were lowest in NSCLC-like cells, rose across cSCLC, and peaked in pure SCLC cells. Taken together, our results capture a coherent molecular trajectory in which immune-interactive, ECM-remodeling NSCLC states progressively relinquish microenvironmental crosstalk and adopt a DDR-high, OXPHOS-addicted, autophagy-supported program characteristic of classical SCLC. The breadth of intermediate states in cSCLC underlines the phenotypic plasticity of cSCLC and suggests therapeutic windows in which targeting metabolic rewiring or exploiting transient immune visibility could intercept lineage switching before full neuroendocrine commitment.

To further evaluate whether cSCLC cells adopt transcriptional profiles representing true intermediates between NSCLC and SCLC, we stratified NSCLC samples by their pathological annotation as LUAD and LUSC and applied an approach complementary to deconvolution. Specifically, we applied quadratic programming^43,44^ assuming that intermediate states would manifest as linear combinations of NSCLC and SCLC profiles. Using pure LUAD, LUSC and SCLC single cells from the reference dataset^24,25^ as endpoints, we observed that bona fide reference cells concentrated at the extremes of the continuum, as expected. In contrast, single cSCLC tumor cells across the cohort exhibited unimodal distributions in the intermediate zones of the respective LUAD–SCLC axes, rather than forming a bimodal distribution (Figure 7G). This pattern suggests that, instead of reflecting only a gradual shift in tumor composition in which LUAD-like cells are progressively replaced by SCLC-like cells, individual tumor cells in SCLC/LUAD samples indeed occupy intermediate transitional states along the NE transformation axis, confirming that the transitional states observed spatially in P4 and P6 are indeed intermediate/hybrid states rather than a simple mixture of two fixed populations. Interestingly, in cSCLC samples with SCLC/LUSC subtype, we observed a similar pattern, with tumor cells again exhibiting unimodal rather than bimodal distributions in the intermediate zones along the LUSC–SCLC axis (Figure 7G), indicating the presence of hybrid states in these cases and suggesting that plastic transitions along the NE transformation axis might also occur in SCLC/LUSC subtypes, similar to SCLC/LUAD. Collectively, these findings support the hypothesis that individual tumor cells of cSCLC undergo NE transformation through intermediate/hybrid states before fully acquiring an SCLC phenotype.

## Discussion

cSCLC is currently classified as a rare type of SCLC and has been estimated at approximately 2–5% of all SCLC cases in the literature over the past two decades.^2,3^ The diagnosis of cSCLC is primarily established on pathological evaluation of surgically resected tumor specimens,^2-6^ and this may cause underdiagnosis in non-surgical patients because accurate diagnosis of cSCLC through small-sized biopsies is challenging. Meanwhile, cSCLC as a subtype of SCLC is currently treated as pure SCLC without specialized treatment despite containing NSCLC (or LCNEC) components, which could be attributed to unclear clonal origin and molecular characteristics of morphologically distinct components within cSCLCs. In this study, we employed spatially resolved multi-omics methods to thoroughly investigate genomic and transcriptomic characteristics of cSCLC with unprecedented resolution, providing a comprehensive resource for dissecting intratumoral heterogeneity and lineage relationships in this entity. In particular, compared to the literature, the cSCLC cohort involved in this study has the highest proportion of SCLC/LUAD patients (>50%), which enables detailed analysis of LUAD–SCLC transitions and of how LUAD-derived components shape the biology and treatment vulnerabilities of cSCLC.

Multi-region WES has demonstrated that SCLC and NSCLC (or LCNEC) components share a common clonal origin despite their divergent histologic appearances.^14^ Although shared ancestry of these components has been suggested by prior studies,^7-10,14^, our spatially resolved genomic profiling of microdissected regions provides direct evidence that NSCLC-specific driver mutations (for example, *EGFR*, *KRAS*, *BRAF*, and *PIK3CA*) are truncal events present across both SCLC and NSCLC/LCNEC compartments. This has immediate diagnostic implications: even a small biopsy containing only SCLC morphology, which is a frequent scenario in unresectable cSCLC where limited sampling from SCLC-predominant primaries often fails to capture co-existing NSCLC (or LCNEC) regions, can still reveal NSCLC-type driver mutations that reflect the underlying combined phenotype, helping to mitigate the limitations of small-sized biopsies in cSCLC diagnosis. Building on this rationale, we developed a targeted 4-gene “cSCLC Detector” panel focused on LUAD- and LUSC-specific driver mutations (*EGFR*, *KRAS*, *BRAF*, and *PIK3CA*), enabling molecular detection of cSCLC from small tissue biopsies or liquid biopsy samples. To maximize diagnostic specificity and facilitate clinical translation, the panel was deliberately restricted to canonical, lineage-informative driver mutations, without incorporating gene fusions or less specific alterations that are more challenging to assay robustly and are less discriminating for a cSCLC-like genotype. In an independent biopsy cohort and a blood ctDNA cohort, this assay identified cSCLC-like cases at rates of 14.1% and 14.2%, respectively, compared with the 2–5% prevalence estimated from surgically resected specimens. The detection sensitivity for the SCLC/LUSC subtype is substantially lower than for SCLC/LUAD, due to the lack of highly specific and recurrent point mutations in LUSC. This could be further improved by identifying additional highly prevalent, LUSC-enriched genomic alterations for ctDNA-based inference and by incorporating orthogonal ctDNA features, such as methylation patterns and fragmentomic signatures, that are compatible with liquid biopsy workflows.^45^ While histopathological confirmation from tumor tissue remains the standard for definitive cSCLC diagnosis^2-6^, the truncal distribution of NSCLC-type driver mutations across components, together with the performance of the cSCLC Detector, suggests that cSCLC is likely underrecognized in routine practice and underscores the clinical need to improve its detection. These observations also support the clinical plausibility that the cSCLC Detector can molecularly flag otherwise unrecognized cSCLC in surgically unresectable patients, although its use as a screening or diagnostic assay will require rigorous validation in prospective clinical studies.

Across these spatial and single-cell analyses, cSCLC emerges as a tumor type in which histologically distinct domains are embedded in sharply contrasting microenvironmental niches. NSCLC/LCNEC regions support an “inflamed” TME with infiltrated T cells, macrophages, and inflammatory iCAFs, whereas adjacent SCLC regions are relatively immune-desert and show active exclusion of infiltrating immune and stromal cells. At the transcriptional level, SCLC-enriched domains consistently exhibit reduced expression of HLA class I genes and *B2M* compared with neighboring NSCLC/LCNEC regions, pointing to lineage-associated downregulation of antigen presentation as one mechanism of impaired T cell engagement. In parallel, fibroblast-rich bands at the interface of these domains are consistently enriched for the CAF S3 program and display a COL11A1⁺, ECM-remodeling myCAF-like signature, closely resembling the COL11A1⁺ boundary CAFs recently implicated in collagen-mediated T cell exclusion and immunotherapy resistance in NSCLC.^26^ In cSCLC, these CAF S3/COL11A1⁺ domains form collagen-dense stromal “septa” that segregate SCLC from NSCLC/LCNEC compartments and coincide with lymphocyte-poor SCLC territories, suggesting that CAF-driven matrix remodeling and antigen-presentation downregulation act in concert to sustain an immune-excluded SCLC niche within otherwise immune-infiltrated tumors. Although our cohort lacks outcome data to directly link these features to prognosis or treatment response, the convergence of MHC-I suppression and CAF S3/COL11A1⁺ barrier architecture, together with prior data in NSCLC and pure SCLC, raises the testable hypothesis that stromal and lineage-specific immune-evasion programs may uncouple the presence of immune infiltrates from actual sensitivity to immune checkpoint blockade in cSCLC, a possibility that warrants prospective clinical and spatially resolved correlative studies. More broadly, this composite TME, characterized by immune-hot NSCLC/LCNEC domains juxtaposed with immune-cold, antigen-presentation-low SCLC domains encased by CAF S3-rich stroma, underscores the need to consider regional context when designing therapies and suggests that combining lineage-informed treatment with strategies that restore antigen presentation and/or modulate pro-tumor CAFs or collagen architecture may be required to fully engage antitumor immunity in cSCLC.

A fundamental question regarding cSCLC pathogenesis is whether the mixed histology arises from the passive outgrowth of pre-existing, distinct states resembling “pure” SCLC or NSCLC, or through active lineage plasticity in which tumor cells adopt intermediate phenotypes. Prior bulk-level work, such as the study by Quintanal-Villalonga and colleagues,^13^ used bulk multi-omics of pre- and post-transformation specimens to infer similar LUAD-to-SCLC NE transformation trajectories and proposed a model in which NE-transformed SCLC arises from a pre-existing LUAD clone with a permissive genomic context, rather than being present as a large, fully transformed SCLC component at the outset. However, bulk averaging could not definitively distinguish whether the transformation arose from gradual reshuffling of coexisting clones or from transitioning hybrid cells. By anchoring our ST and snRNA-seq data to single-cell reference atlases of LUAD, LUSC, and canonical SCLC-A, -N, and -P subtypes,^24,25^ we found that cSCLC tumor cells populate a continuous NSCLC-to-SCLC identity axis rather than segregating into two discrete clouds, and deconvolution against pure NSCLC and SCLC endpoints yields unimodal distributions centered at intermediate positions rather than the bimodal pattern expected from a simple mixture of fixed NSCLC and SCLC clones. This supports a model in which cSCLC contains tumor cells that reside in intermediate, transcriptionally hybrid states rather than representing a mere collage or gradual reshuffling of preexisting NSCLC and SCLC populations. Within the SCLC-like compartment, mapping onto A/N/P subtype space reveals a similar theme: whereas pure SCLC cells cluster tightly near single-subtype vertices, roughly one-third of SCLC-like cells in cSCLC occupy hybrid regions with concurrent activation of A, N, and/or P programs, including NE–POU2F3 combinations (P14) that are rare in *de novo* SCLC, indicating markedly increased subtype transcriptional plasticity compared to pure SCLC.^33^ Together, these observations argue that the transcriptional signatures in cSCLC are not solely due to regional admixture of distinct subtype-specific cells but reflect genuine single-cell plasticity along both the NSCLC-to-SCLC transformation axis and the A/N/P subtype continuum.

Our *EGFR* data provide further indirect support for a LUAD-first transformation model. In cSCLC with the SCLC/LUAD subtype, *EGFR* mutations are present in 40% of patients and are detected across essentially all sampled SCLC and LUAD regions, indicating that *EGFR* is a truncal event shared by both components. The mutation spectrum (exon 19 deletions, L858R, E709V/G719C) and this 40% frequency closely match that reported for Asian LUAD patients, while being much higher than the ∼3% frequency seen in pure *de novo* SCLC. If cSCLC primarily arose from a pre-existing SCLC that secondarily acquired a minor LUAD component, one would not expect a LUAD-like truncal *EGFR* mutation frequency and clonal distribution. Instead, these patterns are more consistent with LUAD-like, *EGFR*-driven clones that later undergo NE transformation into SCLC, consistent with prior reports.^13^ Nonetheless, it remains inherently difficult to fully exclude the presence of transformed cells at very early stages when working with clinical specimens, particularly in untreated cases, which are typically collected at a single time point. Tracking such rare events will require multi-region longitudinal sampling and/or single-cell lineage tracing and should be a priority for future work.

Our findings also complement and extend prior work on LUAD-to-SCLC transformation, particularly the multiomic analysis by Quintanal-Villalonga and colleagues.^13^ That study used bulk exome and RNA sequencing of LUAD tumors that had undergone neuroendocrine transformation to SCLC, including combined LUAD/SCLC specimens in which the LUAD (T-LUAD) and SCLC (T-SCLC) components were microdissected and analyzed to infer lineage trajectories and nominate molecular drivers. Our spatial WES results yield consistent conclusion, and both studies support that the best-supported risk factors for predicting the NE transformation are the combination of *TP53*/*RB1* loss and 3p arm loss, although definitive predictive biomarkers will require prospective analysis of statistically sufficient LUAD biopsy samples collected prior to NE transformation. However, bulk averaging could not resolve spatial architecture, tumor–stroma organization, or whether “mixed” signatures reflected coexisting clones or hybrid cells. Our work directly visualizes the juxtaposition of SCLC and NSCLC/LCNEC domains, defines their immune and fibroblast niches, and reveals that tumor cells in cSCLC occupy intermediate positions along the NSCLC-to-SCLC NE differentiation axis. We demonstrate that immune-excluded SCLC regions couple MHC-I downregulation with CAF S3/COL11A1⁺ boundary programs, and identify extensive hybrid A/N/P subtype states that cannot be explained by simple regional admixture. Moreover, by extending the analysis beyond SCLC/LUAD to include SCLC/LUSC and SCLC/LCNEC cases and demonstrating shared truncal drivers across components in each, we generalize key principles of lineage plasticity and TME reprogramming beyond the LUAD-only setting examined previously and provide a spatially resolved, single-cell view of how mixed histology is organized across distinct histologic subtypes of cSCLC. Collectively, our results provide a foundation for understanding cSCLC evolution and advancing innovative diagnostics and therapeutics.

### Limitations of the study

Our study provides a spatially resolved multi-omics characterization of cSCLC. However, several limitations should be considered when interpreting the results. First, the cohort size is relatively small (n = 19), reflecting both the rarity of cSCLC and the practical challenges of obtaining high-quality multi-region specimens with matched multi-omics data. Although this represents one of the largest molecularly profiled cSCLC cohorts reported to date, the limited sample number restricts statistical power and may limit the detection of subtype-specific patterns or rare molecular events. Second, the histological subtype distribution within the cohort was uneven, with SCLC/LUAD cases comprising more than half of the samples. This imbalance may affect the generalizability of our findings to other cSCLC subtypes, such as SCLC/LUSC or SCLC/LCNEC, which were less represented. Future studies with larger, more balanced cohorts and longitudinal sampling will be required to validate these findings, establish robust biomarkers, and clarify the clinical relevance of tumor plasticity and microenvironmental niches identified here.

## Acknowledgements.

We thank the following agencies and foundations for support: National Key Research and Development Program of China (2023ZD0501905 to Z.L.), National Natural Science Foundation of China (82473288 to Z.L., 22374027 to Q.S., 82202626 to Z.W.), NIH U54CA274509 (to W.W.), Andy Hill CARE Fund (FY23-BSF-01 to W.W.). Program of Shanghai Academic/Technology Research Leader from Shanghai Science and Technology Committee (23XD1403900 to Q.S.), Shanghai Municipal Health Commission (2022XD029 to Z.L.), Beijing Xisike Clinical Oncology Research Foundation (Y-2024AZ(EGFR)ZD-0260 to Z.L.), Yangtze River Delta Joint Sci-Tech Innovation and Research Projects (2023CSJZN0600 to Z.L.).

## AUTHOR CONTRIBUTIONS

S.L., W.W., Q.S. and Z.L. conceived the project. Z.W., L.L., W.D., Y.Z., L.Z., X.O., and W.X. performed the experiments. Z.W., Q.L., J.W., R.Q., W.W., and Q.S. analyzed the data. J.W., Y.Y., and Z.L. collected the clinical specimens and analyzed the clinical data. Z.W., Q.L., W.W., and Q.S. wrote the manuscript. All authors contributed to writing the paper and discussing the content and approved the final manuscript.

## KEY RESOURCES TABLE STUDY DESIGN AND PATIENT INFORMATION

### Human patient samples

This study was conducted at Shanghai Chest Hospital and enrolled 19 treatment naive cSCLC patients from 2011 to 2018 for multi-omics profiling, including multi-region WES, ST and snRNA-seq. This study was performed according to the principles of the Helsinki Declaration and was approved by the institutional review board (#IS2117). Written informed consent has been obtained from all participants. The SCLC and NSCLC regions of formalin-fixed and paraffin-embedded (FFPE) cSCLC samples were defined by experienced pathologists based on H&E review, supported by a panel of IHC markers covering canonical SCLC and NSCLC (or LCNEC) markers used routinely in clinical diagnosis (including CD56, chromogranin A (ChgA), synaptophysin (Syn), TTF-1, Napsin A, p40, CK5/6, and CK7). Clinical information for the patients is provided in Table S1. Sex was assigned (18 male and 1 female patients). Sex-based analyses were not performed due to the limited sample size. Gender was not determined.

| No. | Subtype | WES (# of regions) | ST | snRNA-seq |
| --- | --- | --- | --- | --- |
| P1 | SCLC/LUAD | Yes | / | / |
| P2 | SCLC/LUAD | Yes | / | / |
| P3 | SCLC/LUAD | Yes | / | Yes |
| P4 | SCLC/LUAD | Yes | Yes | Yes |
| P5 | SCLC/LUAD | Yes | / | / |
| P6 | SCLC/LUAD | Yes | Yes | Yes |
| P7 | SCLC/LUAD | Yes | / | / |
| P8 | SCLC/LUAD | Yes | / | / |
| P9 | SCLC/LUAD | Yes | / | / |
| P10 | SCLC/LUAD | Yes | / | / |
| P11 | SCLC/LUSC | Yes | Yes | Yes |
| P12 | SCLC/LUSC | Yes | / | Yes |
| P13 | SCLC/LUSC | / | / | Yes |
| P14 | SCLC/LCNEC | Yes | Yes | Yes |
| P15 | SCLC/LCNEC | Yes | Yes | Yes |
| P16 | SCLC/LCNEC | Yes | / | Yes |
| P17 | SCLC/LCNEC | Yes | / | Yes |
| P18 | SCLC/LCNEC | / | Yes | Yes |
| P19 | SCLC/LCNEC | / | / | Yes |

## METHOD DETAILS

### Laser capture microdissection (LCM)

For multi-region WES of 16 patients, LCM (Molecular Machines & Industries, MMI Cellcut Plus) was used to isolate SCLC and NSCLC or LCNEC regions of interest from FFPE cSCLC tissue sections based on H&E staining and pathological evaluation. For LCM, five patients had one tumor block (P1, P2, P13, P14 and P15), and 8 patients had two independent tumor blocks (P3, P4, P5, P6, P7, P8, P9 and P12), and 3 patients had three independent tumor blocks (P10, P11 and P16). The microdissected tumor samples were collected in 1.5 mL tubes for DNA isolation.

### Genomic DNA isolation and whole exome sequencing (WES)

Genomic DNA was isolated using the GeneRead^TM^ DNA FFPE Kit (Qiagen) from LCM-isolated, FFPE tumor samples according to the GeneRead^TM^ DNA FFPE Handbook. The DNA concentration was measured using Qubit 4.0 Fluorometer (Invitrogen) with Qubit dsDNA HS (High Sensitivity) Assay Kit (Invitrogen). The DNA isolated from FFPE tumor samples and matched normal tissues were subjected to WES. A total of 200 ng of genomic DNA was sheared into 200-250 bp fragments with Covaris S2 ultrasonicator (Covaris). Sequencing libraries were prepared by using the KAPA DNA Library Preparation Kit (Kapa Biosystems) according to the manufacturer’s protocol. Briefly, the ends of the fragments were repaired before "A" addition, followed by adapter ligation, amplification, and hybridization to the SeqCap EZ library. Exome regions were captured with SureSelectHuman All Exon V6 (Agilent Technologies) according to the manufacturer’s protocol. Recovered libraries were amplified by PCR. Samples with DNA library concentration ≥20 ng/uL were sequenced with 100 bp paired-end on a Geneplus-2000 sequencing platform (Geneplus, Beijing, China).

### Isolation of nuclei from snap-frozen human lung cancer tissues

Cell nuclei were extracted from snap-frozen tissues using GEXSCOPE^®^ Single Nucleus RNA Library Kit V2 (Singleron Biotechnologies). On ice, the tissue was immersed in a cold nucleus separation solution and cut into small pieces. Kimble douncer (KIMBLE® KONTES^®^ Dounce Tissue Grinder) was employed to homogenize snap-frozen tissues, followed by on-ice incubation for 15 min. After homogenization and digestion, the nuclei suspension was filtered with a 40 µm cell strainer and centrifuged at 200× g for 2 min at 4℃. The separated supernatant was centrifuged at 500× g for 5 min at 4℃. The nuclei pellet was resuspended in 0.25 ml of cold nuclei suspension buffer. The quality of nuclei was assessed by Trypan Blue staining (0.4% w/v, Invitrogen) under a light microscope. The concentration of single nucleus suspension was adjusted to 1 × 10^5^ cells/ml in PBS for subsequent library preparation.

### Spatial transcriptomics (ST) sequencing of snap-frozen tumor tissues

Sections of snap-frozen tumor tissues were cut into 10 μm thickness and mounted onto the GEX arrays. Cryosectioned tissue samples were placed on a Thermocycler Adaptor with the active surface facing up and incubated at 37℃ for 1 min, followed by fixing with methyl alcohol at -20℃ for 30 min and H&E staining. The H&E stained images were recorded by an Olympus SLIDEVIEW VS200 slide scanner. The permeabilization time of cryosections on the slides was determined using Visium Spatial Tissue Optimization Slide & Reagent Kit according to the manufacturer’s introductions. Visium spatial libraries were constructed using Visium Spatial Gene Expression Slide & Reagent Kit (10× Genomics) according to the manufacturer’s introduction. Slide cassette was used to create leak-proof wells for adding reagents. Approximately 70 μL of permeabilization enzyme was added into the wells and incubated at 37℃ for 15min. Each well was washed with 100 μL of 1×SSC, followed by adding 75 μL of reverse transcription master mix for cDNA synthesis. At the end of first-strand cDNA synthesis, the cDNA synthesis reaction buffer was removed from the wells and 75 μL of 0.08 M KOH solution was added for incubation at room temperature for 5 min. After removal of KOH solution, the wells were washed with 100 μL of EB buffer, and 75 μL of second strand master mix was added to each well for second-strand synthesis. cDNA amplification was performed on a Thermal Cycler (Bio-Rad). Spatial gene expression library was constructed using amplified cDNA and the libraries were sequenced on an Illumina NovaSeq6000 sequencer with a sequencing depth of at least 100,000 reads per spot with a paired-end 150 bp (PE150).

In this study, we performed ST on frozen tissue sections from 6 cSCLC patients, including 2 SCLC/LUAD, 1 SCLC/LUSC and 3 SCLC/LCNEC patients. We obtained a median of 1,573 voxels (range: 1,023∼2,556) in each sample with a median of 20,417 (range: 20,236∼21,913) genes.

### Single nucleus RNA-Seq (snRNA-seq) library preparation and sequencing

Single nucleus suspension was loaded onto the microfluidic chip (GEXSCOPE^®^ Single Nucleus RNA Library Kit V2, Singleron Biotechnologies) for snRNA-seq library preparation according to the manufacturer’s instructions. Quality control for amplified cDNA was conducted by Qsep100 (Bioptic). The generated snRNA-seq libraries were subjected to sequencing on an Illumina NovaSeq 6000 instrument using a paired-end 150 bp approach. In this study, snRNA-seq was performed on frozen tissue samples from 12 cSCLC patients. We obtained a median of 7,268 cells (range: 2,966-26,745) measured per sample and a total of 107,842 cells with 19,786 genes detected.

### Multiplex immunofluorescence staining

Formalin-fixed paraffin-embedded (FFPE) tissue sections were cut into 5 μm sections and mounted on glass slides. Sections were baked at 62°C for 1 h, followed by deparaffinization in fresh xylene (3 × 20 min) and then rehydrated in 100% (2 × 1 min), 95% (2 × 1 min), and 70% (1 min) alcohol in sequence. Slides were rinsed in running tap water for 5 min, and briefly washed in distilled water. Endogenous peroxidase activity was quenched by incubating sections with 3% H_2_O_2_ for 10 min at room temperature, followed by phosphate-buffered saline (PBS) washes (3 × 5 min). Heat-induced epitope retrieval was performed in 1 mM Tris–EDTA buffer (pH 9.0) by boiling for 15 min and maintaining at temperature for an additional 15 min, then allowing slides to cool naturally to room temperature. After PBS washes (3 × 5 min), sections were blocked with 5% bovine serum albumin (BSA) for 20 min at room temperature and incubated with the first primary antibody (Syn, Napsin A, PDGFRβ, or CD8; diluted in antibody diluent) overnight at 4°C. Slides were equilibrated for 1 h at room temperature (or 30 min at 37°C), washed in PBS (3 × 5 min), and incubated with an HRP-conjugated secondary antibody (goat anti-rabbit IgG-HRP, 1:2000) for 30 min at 37°C. TSA signal amplification was then performed using tyramide working solutions (Try-488, Try-594, Try-Cy3, or Try-Cy5; ready-to-use) for 10-30 min at room temperature, followed by PBS washes (3×5min). For multiplex staining, antibody stripping/epitope re-retrieval was achieved by repeating the heat retrieval step in Tris–EDTA (pH 9.0), after which the next primary antibody was applied and the HRP/TSA deposition cycle was repeated using spectrally distinct fluorophores. After completion of multiplex staining, nuclei were counterstained and sections were mounted with an anti-fade mounting medium containing DAPI.

### Immunohistochemistry (IHC)

Immunohistochemistry was performed on formalin-fixed, paraffin-embedded (FFPE) tissue sections (4 μm thick). Slides were baked at 60°C for 2 hours, deparaffinized in xylene, and rehydrated through a graded ethanol series (100%, 90%, 70%, and 50%) to PBS. Endogenous peroxidase activity was quenched by incubation with 3% hydrogen peroxide (H2O2) for 15 min. Heat-induced epitope retrieval (HIER) was performed using sodium citrate buffer. The sections were then incubated with primary antibodies against MASH1 (1:200; Abcam, Cat# ab211327), NeuroD1 (1:200; Abcam, Cat# ab205300), and POU2F3 (1:200; Cell Signaling Technology, Cat# 36135S). Subsequently, the tissues were incubated with a biotin-conjugated secondary antibody, followed by the avidin-biotin-peroxidase complex. Staining was visualized using diaminobenzidine (DAB), and sections were mounted with neutral balsam. IHC staining results were evaluated by experienced pathologists who were blinded to the clinical data. The expression status was classified into three categories based on the staining intensity and distribution: Positive, defined as strong protein expression; Partial Positive, defined as focal, scattered, or patchy positivity; and Negative, defined as the absence of protein expression.

### Mutation calling in WES data and construction of phylogenetic trees

The raw genomic data generated from WES were aligned to human genome hg19 (UCSC) using BWA (v0.7.10)^46^ with default parameters. SAM files were converted to BAM files and sorted by chromosomal coordinates using Samtools (v1.98)^47^. Picard (v1.56) was used to mark PCR duplicates. Realignment and recalibration of sorted and marked BAM files were performed by GATK (v4.0)^48^. Single nucleotide variants (SNVs), small insertions and deletions (Indels) were called by MuTect (v1.1.4). All variants were annotated by SnpEff 4.0. Filtering criteria were then applied to identify candidate somatic mutations when (i) the mutation was detected in at least 5 high-quality reads; and (ii) the mutation had a variant allele frequency >0.01; and (iii) the mutation was not present in>1% of the population in the ExAC, 1000G and gnomAD databases; and (iv) the mutation was absent from the normal samples. All somatic mutations identified in the samples were summarized in Table S3. Somatic copy number alterations (CNAs) were identified with GATK (v4.0).^48^

Phylogenetic relationships were reconstructed using nonsynonymous somatic mutations identified from multi-region whole-exome sequencing of laser-capture microdissected tumor regions. For each patient, regions obtained from distinct histological areas and tumor blocks were treated as separate evolutionary units. A binary mutation matrix was constructed indicating the presence or absence of each nonsynonymous mutation across regions within the same tumor. Pairwise distances between regions were computed based on their mutational profiles, and phylogenetic trees were inferred using the neighbor-joining method implemented in the ape R package.^49^

### 10x Visium spatial RNA-seq data preprocessing

We processed Visium spatial RNA-seq data and brightfield microscope images using the Space Ranger pipeline (v1.0.0; 10X Genomics). This pipeline performed tissue detection, STAR-based read alignment, feature-spot matrix generation, clustering, gene expression analysis, and spatial spot placement. Subsequent analysis of the unique molecular identifier (UMI) count matrix utilized the Seurat R package (v4.1.1).^50^ Data normalization was first conducted via SCTransform^51^ to correct for sequencing depth variation and identify high-variance features, with results stored in the SCT assay. The gene expression matrix was then aligned with the positional information from the H&E image.

### Integrative clustering and annotation of ST spots

To explore the cellular characteristics of spatial spots across different samples, we integrated all ST data samples into a unified dataset. We then co-embedded all ST spots into a UMAP space using Seurat and performed clustering to identify groups of spatial spots with similar transcriptomic profiles.

For cluster annotation, we leveraged a carefully curated reference dataset^24,25^ containing annotations for NSCLC or LCNEC and SCLC subtypes, as well as various immune and stromal cell types, including T cells, B cells, plasma cells, mast cells, macrophages, neutrophils, and fibroblasts. To assign clusters, we applied a combined similarity metric that equally weighted Jensen-Shannon divergence and cosine similarity. Jensen-Shannon divergence quantifies differences in probability distributions, making it effective for distinguishing gene expression patterns across clusters, while cosine similarity captures the directional similarity between expression vectors, reflecting global transcriptional trends. By integrating these two metrics, we leveraged both distributional and vector-based similarities, ensuring a robust and reliable comparison between spatial clusters and the reference single-cell RNA-seq dataset.^24,25^ These metrics quantified the resemblance of spatial clusters to predefined cell types.

Finally, to confirm the accuracy of cluster annotations, we examined the expression of known marker genes for specific cell types, reinforcing the biological relevance of the identified clusters.

### Single-cell deconvolution of ST

To further resolve the cellular composition of individual spatial spots, we employed the SPOTlight algorithm.^30^ This algorithm deconvolutes the transcriptomic profile of each spot into probabilistic contributions from predefined single-cell clusters. To ensure its ability to differentiate closely related cell types, such as SCLC subtypes, we used distinguishing marker genes for each cell type. This was achieved through pairwise differential expression analyses using the reference dataset.^24,25^ For instance, to determine genes highly expressed in SCLC-A, we recursively compared SCLC-A to SCLC-N, SCLC-P, and all other cell types. This analysis was conducted using Seurat’s FindMarkers function with "only.pos = TRUE", ensuring that only upregulated genes in SCLC-A were retained. Next, we refined the selection by filtering for genes with a fold change greater than 1.5 across all comparisons. The intersected set of these genes represented the uniquely and highly expressed markers specific to SCLC-A. We repeated this process for all cell types, ensuring that each was characterized by a distinct set of marker genes that effectively differentiated it from others in the reference dataset.^24,25^ The resulting marker genes then served as the foundation for comparing spatial clusters with reference cell types.

To improve the specificity of deconvolution, we further intersected the marker genes with the top 5000 most variable genes from the ST dataset. This step prioritized spatially informative genes, reducing noise from lowly expressed or uninformative features and enhancing the alignment between single-cell and spatial data. Additionally, we tailored the NSCLC component of the reference dataset based on the diagnostic type of each sample. Specifically, for samples diagnosed as LUAD, LUSC, or LCNEC combined with SCLC, we only included the corresponding non-small cell subtype (LUAD, LUSC, or LCNEC) alongside SCLC, while retaining all immune and stromal cell types in the reference. This ensured that the deconvolution process remained biologically relevant to the specific tumor composition of each sample without excluding critical immune or stromal components.

By leveraging the same reference dataset used for cluster annotation, SPOTlight^30^ provided relative proportions of different cell types or states within each spatial spot, ensuring consistency in cell type identification across analyses.

### Generation of single-nucleus gene expression matrices

For snRNA-seq, raw reads were processed to generate gene expression matrices by CeleScope™ (v1.14.1). Briefly, low-quality reads, adapter sequences and polyA tails were removed by Cutadapt (v4.6)^.52^ The trimmed Read2 reads were mapped to the reference genome GRCh38 (Ensembl version 92 annotation) via STAR.^53^ Reads were grouped together to count the number of UMIs per gene per cell. For quality control, we excluded cells with a unique feature count below 1000 or cells with mitochondrial counts ratio greater than 10%.

### snRNA-seq data analysis

Quality control and preprocessing of cSCLC snRNA-seq data were performed using Seurat (version 5). Nuclei with fewer than 200 detected genes or with a mitochondrial gene fraction exceeding 10% were excluded from downstream analysis. We integrated the cSCLC data with two well-annotated public datasets. First, we identified 4,000 highly variable genes (HVGs) across the combined dataset, following Seurat’s standard workflow. Principal component analysis (PCA) was performed on the scaled expression matrix using Seurat’s default parameters. The top 15 principal components (PCs) were then used as input for batch correction with Harmony, which was run with default parameters unless otherwise specified.

To validate that batch correction effectively removed technical batch effects without causing overcorrection, we examined the clustering patterns of biologically related and distinct cell types. We found that previously annotated NSCLC cells from two independent reference datasets^24,25^ aligned well with each other in the integrated space, positioning closely but remaining clearly distinguished from SCLC cells. Additionally, shared cell types such as immune cells (e.g., T cells, macrophages) and fibroblasts clustered together across datasets, demonstrating successful batch alignment and resolution of major cell types within the integrated reference. These immune and fibroblast populations were well separated from tumor cells overall, reflecting preservation of expected biological relationships. Notably, we observed some cSCLC cells overlapping with immune and fibroblast clusters, and these overlapping cSCLC cells expressed marker genes characteristic of immune or stromal lineages, suggesting the presence of non-tumor nuclei in cSCLC samples or potential tumor–stromal interactions.

For cell type annotation in cSCLC, we assigned identities by examining where cSCLC nuclei clustered relative to the resolved reference cell types in the integrated dataset. Specifically, cSCLC nuclei were annotated based on the reference populations (e.g., NSCLC tumor cells, SCLC tumor cells, immune cells, fibroblasts) with which they clustered most closely, supported by the expression of canonical marker genes indicative of each lineage. The majority of cSCLC nuclei, however, formed distinct clusters that stood out from both NSCLC and SCLC reference populations. These cSCLC-specific clusters expressed canonical SCLC markers, including ASCL1, supporting their classification as SCLC-like tumor cells. To further confirm the malignant nature of these populations, we reinforced their tumor identity through copy number alteration (CNA) analysis, as detailed in the next section.

### Copy Number Alteration analysis

To validate our tumor region identification in ST and tumor cell annotations in snRNA-seq, we applied InferCNV^54^ to both, allowing us to infer large-scale chromosomal copy number alteration (CNA), a hallmark of tumor cells.

For ST data, we pooled all spatial transcriptomics samples together and designated Cluster 1, which primarily consisted of fibroblasts and immune cells, as the reference population. To further refine the reference, we ensured minimal tumor contamination by removing any spots where the inferred tumor component from deconvolution analysis exceeded 0.1. This allowed us to establish a genomically stable baseline for CNA inference.

For snRNA-seq data, we performed InferCNV on an integrated dataset that included our snRNA-seq dataset and the reference datasets.^24,25^ To ensure computational efficiency and maintain statistical power, we downsampled the integrated dataset to 30,000 cells before running InferCNV. For reference cells, we used annotated immune cells from the reference dataset, assuming they represented a genomically stable population.

Both analyses were conducted using the InferCNV package in R, with the following key parameters: hmm = TRUE (enabling Hidden Markov Model for CNA calling), denoise = TRUE (to reduce noise in CNA signal), noise cutoff = 0.1, gene expression percentage cutoff = 0.01. Other parameters were set to default values as recommended by InferCNV. To identify tumor cells based on inferred CNA burden, we established a cutoff using a two-step approach. First, we inspected the histogram of CNA burden scores (Figure 7C) and identified an inflection point, a point where the distribution shifted from a sharp decline (representing normal cells) to a broader distribution with elevated CNA levels (indicative of tumor cells). Second, we validated this tentative cutoff by cross-referencing it with known normal and tumor cell populations in our dataset to ensure that the chosen threshold accurately distinguished these groups. Based on this analysis, we selected a CNA burden cutoff of approximately 0.04 (as indicated by the dashed line in the histogram) to classify cells as tumor or normal.

These analyses allowed us to confirm tumor cell identity and tumor-associated regions in both spatial transcriptomics and single-nucleus RNA-seq datasets, strengthening the reliability of our annotations.

### Mapping spatial patterns

To analyze the spatial organization and relationships between cell types, we employed Ripley’s K function, a statistical method for assessing spatial point patterns. This function compares the observed spatial distribution of points (e.g., cell types or components) to a theoretical random distribution, known as complete spatial randomness (CSR), across increasing spatial scales. K values above the expected CSR values indicate clustering, suggesting spatial association, while values below indicate dispersion, reflecting a more scattered distribution.

We performed this analysis in R using the spatstat package, specifically the Kest function, which computes Ripley’s K function for a given set of spatial point coordinates. Before applying the Kest function, we formatted the spatial spot coordinates as a point pattern object (ppp) using geom::pp function, allowing us to analyze spatial clustering patterns across increasing distances.

To ensure the inclusion of spatial spots with a meaningful representation of each cell type, we determined a threshold for the deconvolution score based on the third quartile (Q3) of the scores for each cell type or group. If Q3 exceeded 0.3, we set 0.3 as the threshold; otherwise, we used Q3. This process was implemented in R using the quantile() function to compute Q3 for each cell type’s deconvolution scores. We then filtered spatial spots based on this threshold before applying the Ripley’s K function analysis.

To visualize the results, we plotted K(r) as a function of distance (r) for different cell type pairs across multiple patient samples. These plots allow us to assess the degree of clustering or dispersion between NSCLC and SCLC components, as well as their interactions with fibroblasts and immune cells. Specifically, different colored lines represent distinct cell type pairwise comparisons, with the background (CSR) shown as a dotted line. Elevated K values above the background indicate clustering, suggesting that the two cell types are frequently located near each other, while lower K values suggest spatial segregation.

This analysis provides insights into how different tumor and immune cell types are spatially organized within the tumor microenvironment, revealing potential interactions, co-localization patterns, or exclusion dynamics that may be biologically significant.

### Gene program scoring and functional annotation

The activity of the gene program across individual cells was profiled by snRNA-seq and across spatial spots in Visium data using AUCell.^37^ AUCell quantifies gene set activity by ranking genes within each cell or spot according to expression and calculating the area under the recovery curve for genes in the gene set among the highest-ranked genes, thereby yielding a robust activity score for each cell or spot. We scored each cell or spot using the defined gene set. To interpret the biological processes represented by a particular gene set or list, we performed Gene Ontology enrichment analysis using the clusterProfiler R package.^55^ GO terms were tested for over-representation against a background of all detected genes in the dataset, and statistical significance was assessed using a hypergeometric test with false discovery rate correction. The top ten enriched GO terms, ranked by adjusted P value, were selected for visualization. The S3 CAF gene set used for Figures 6F and 6G was derived from the S3 CAF signature genes as reported.^27^ All other gene sets or gene lists used in this manuscript are described in the corresponding sections below, together with details of how they were generated for each analysis.

### SCLC A, N, P subtype plasticity

In our ST data, we detected co-expression of *ASCL1* and *NEUROD1*, as well as simultaneous expression of *ASCL1* with P-type marker genes. These findings raised the intriguing possibility that cSCLC might display increased plasticity among the A, N, and P subtypes. However, because ST lacks single-cell resolution, we turned to snRNA-seq to investigate this phenomenon at a higher resolution within SCLC-like cells identified in our cSCLC dataset.

First, we performed SPOTlight^30^ deconvolution on the cSCLC snRNA-seq data, using reference profiles from SCLC and NSCLC to estimate cell identities. In this analysis, a score of zero indicated no similarity to a reference cell type, while a score of one denoted complete identity. We classified cells with SCLC identity scores exceeding 0.75 and NSCLC scores below 0.25 as SCLC-like cells, allowing us to isolate a population most representative of the SCLC phenotype within cSCLC.

Next, we sought to determine whether these SCLC-like cells exhibited substantial plasticity across A, N, and P subtypes. We assessed this by testing whether individual SCLC-like cells simultaneously expressed multiple subtype-specific gene expression signatures. The gene sets defining A, N, and P subtypes were curated based on the marker genes identified in earlier sections of our study. To quantify gene set activity, we applied the AUCell package^37^ in R.

To establish a reference baseline, we first performed this analysis on canonical single cells representing each A, N, and P subtype, thereby defining the expected transcriptional profiles for these subtypes. We then applied the same approach to the SCLC-like cells from cSCLC to investigate their distribution across the A-N-P transcriptional space. This strategy enabled us to evaluate whether cSCLC cells demonstrate heightened plasticity by exhibiting mixed subtype signatures or transitions between A, N, and P states.

For visualization, we normalized the AUC scores for A, N, and P within each cell so that their sum equalled one. This normalization allowed each cell’s subtype identity to be represented on a ternary plot, where cells near a vertex displayed predominant expression of a single subtype (A, N, or P), and cells closer to the center reflected a more plastic state characterized by mixed subtype features. We generated ternary plots using the ggtern package in R, overlaying contour density levels to highlight areas with high concentrations of cells.

### Identification of subtype hybrid cells

For each SCLC or SCLC-like (defined previously) cell, we computed a dominance ratio defined as the maximum of the normalized A, N, and P subtype signature scores divided by their mean (max(A, N, P) / mean(A, N, P)), which quantifies the extent to which a single subtype program dominates. Cells belonging to a single subtype are expected to exhibit a high dominance ratio, whereas hybrid cells are expected to show more balanced signature scores and therefore a lower dominance ratio. To establish a classification threshold, we used reference single-subtype (A-, N-, and P-type) SCLC cells as a calibration set. The cutoff was defined as the mean value of max(A, N, P) / mean(A, N, P) across these reference single-subtype cells minus two standard deviations, such that cells falling below this value were classified as hybrid (coANP), whereas cells above the threshold were considered single-subtype–like. This conservative cutoff misclassified only 3.7% of reference single-subtype cells.

### Differential pathway activity analysis

To investigate pathway dynamics across NSCLC, cSCLC, and SCLC cells, we first curated nine functional pathway categories from GO, CP, and HALLMARK gene sets using the msigdbr R package. They included growth and signaling pathways (G1), DNA damage and repair (G2), cell cycle regulation (G3), metabolic pathways (G4), epigenetic regulation (G5), inflammation and immune response (G6), cell death pathways (G7), tumor microenvironment and tissue remodeling (G8), and neural development and signaling (G9). For each pathway, we quantified pathway activity at single-cell resolution by applying the AUCell algorithm^37^ to capture the enrichment of each pathway’s gene set in individual cells. This step produced a matrix of scaled pathway activity scores across all cells.

To establish a unified continuum spanning NSCLC, cSCLC, and SCLC, we used the relative SCLC and NSCLC identity scores previously obtained through SPOTlight^30^ deconvolution on cSCLC samples and extended this approach by performing the same deconvolution on canonical NSCLC and SCLC cells. We normalized the difference between SCLC and NSCLC scores, transforming it into a pseudotime-like trajectory reflecting progression from NSCLC-like to SCLC-like transcriptional states. Next, we employed tradeSeq,^51^ a tool designed to model dynamics along trajectories, to identify pathways exhibiting dynamic changes along this SCLC-NSCLC pseudotime. By fitting generalized additive models, tradeSeq enabled us to detect statistically significant trends in pathway activity associated with transitions between NSCLC and SCLC states. Finally, to visualize these dynamics, we generated a heatmap of scaled pathway activity scores for all pathways across cells ordered by pseudotime, with hierarchical clustering applied to pathways for pattern discovery. Additionally, we overlaid density plots of cell distributions by type (NSCLC, cSCLC, SCLC) along the pseudotime axis to illustrate the positioning of each group relative to the transcriptional continuum.

### Quadratic Programming for intermediate cell state identification

#### (1) LUAD to SCLC transdifferentiation analysis in ST

To investigate whether LUAD to SCLC transdifferentiation would show transcriptionally intermediate states between LUAD and SCLC, and to determine which state our Spatial LUAD and Spatial SCLC samples most closely resemble, we applied quadratic programming (QP) to model their expression profiles as a linear combination of LUAD and SCLC.

We first curated a reference dataset^24,25^ comprising bulk RNA-seq data from four groups: LUAD, T.LUAD (transformed LUAD), T.SCLC (transformed SCLC) and SCLC. To ensure data quality, we performed a Simpson correlation analysis followed by hierarchical clustering to detect and remove implied outlier samples. Then, to put everything on the same scale, we generated *in silico* bulk samples from spatial transcriptomics data, we randomly grouped LUAD and SCLC spatially localized spots from P4 and P6 into three separate *in silico* bulk samples per region. To determine whether T.LUAD, T.SCLC, and the spatial in silico bulk samples exhibit transcriptional profiles indicative of intermediate transitioning states between pure LUAD and pure SCLC, we modelled their gene expression profiles as a weighted combination of LUAD and SCLC. We applied quadratic programming (QP), using LUAD and SCLC bulk RNA-seq profiles as fixed endpoints, and solved for the optimal linear combination that best explains the transcriptional composition of each test sample. This was implemented in R using the quadprog package, specifically the solve.QP() function, which minimizes the squared differences between the observed expression profile and the weighted sum of pure LUAD and pure SCLC expression profiles, subject to the constraint that the weights sum to 1. Finally, we computed the relative contributions of pure LUAD and pure SCLC to each sample, allowing us to infer whether T.LUAD and T.SCLC exhibit transcriptional states that are intermediate between pure LUAD and pure SCLC, and to determine whether the Spatial LUAD and Spatial SCLC in silico bulk samples align more closely with pure LUAD or pure SCLC at the transcriptional level.

#### (2) Intermediate state analysis between LUAD/LUSC and SCLC in cSCLC

For cSCLC samples that were pathologically classified as SCLC mixed with either lung adenocarcinoma (LUAD) or lung squamous cell carcinoma (LUSC), we sought to determine whether these tumors exhibited transcriptional profiles intermediate between LUAD/LUSC and SCLC. To address this, we applied quadratic programming (QP)^43,44^ to model each cSCLC expression profile as a linear combination of LUAD/LUSC and SCLC reference profiles, following an approach similar to that described previously. Briefly, we first established expression profiles for LUAD/LUSC and SCLC by using reference datasets^24,25^ to define the two endpoints representing pure NSCLC and SCLC states. We then used QP to deconvolute the expression profiles of cSCLC samples, estimating their identities as weighted combinations of these endpoints. To quantify each sample’s position along the NSCLC–SCLC continuum, we calculated a final metric by subtracting the LUAD/LUSC identity score from the SCLC identity score. In this metric, a value of 1 indicated a purely SCLC-like transcriptional state, 0 corresponded to a LUAD/LUSC-like profile, and 0.5 represented an intermediate state between the two. Finally, we visualized the distribution of cSCLC samples relative to LUAD/LUSC and SCLC cells by plotting density plots, allowing us to compare the transcriptional positioning of cSCLC against canonical LUAD/LUSC and SCLC populations.

### Tumor microenvironment analysis in multiplex immunofluorescence

Cell phenotypes were defined using a four-marker panel to distinguish tumor lineages and microenvironmental components. Specifically, cells were classified as follows: SCLC tumor cells (Synaptophysin^+^/DAPI^+^), LUAD tumor cells (Napsin A^+^/DAPI^+^), T cells (CD3^+^/DAPI^+^), and cancer-associated fibroblasts (PDGFRβ^+^/DAPI^+^). Whole-slide images were visualized using SliderViewer (v2.6.0), and quantitative spatial analysis was performed using the HALO (v4.1.5944.283) digital pathology platform (Indica Labs). To assess microenvironmental heterogeneity, immune and stromal cell densities were quantified within pathologically annotated SCLC and LUAD regions of the same tumor. Statistical differences in cell infiltration between the two histological components were evaluated using a paired two-sided Mann-Whitney U test.

### Statistical analysis

The normality of the data was tested by the Kolmogorov-Smirnov test. Data are reported as means and SDs, median and interquartile range (IQR), as appropriate. The Mann-Whitney test was performed for the non-parametric test between two groups that were not normally distributed. For analyses involving multiple comparisons, p values were adjusted using the two-stage step-up method of Benjamini, Kriger, and Yekutieli. All statistical analyses were performed with GraphPad Prism 10.

## Supplemental information

### Supplementary Figures

**Figure S1.**
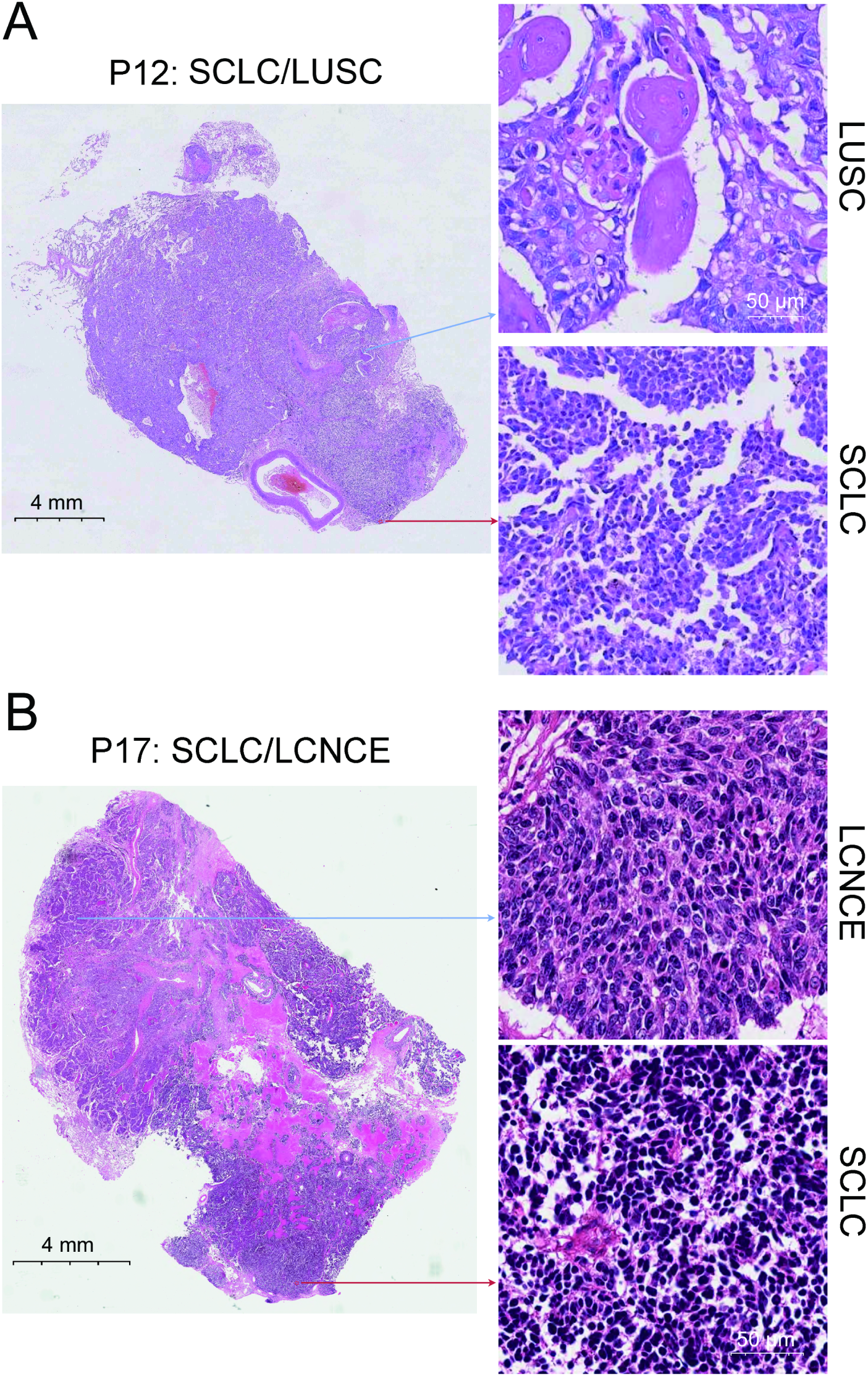
Illustrative FFPE images of two of our cSCLC samples (P12, P17), showing a mixture of SCLC and LUSC or LCNCE morphologic components within the same tumor tissue samples. (A) Coexistence of SCLC and LUSC components within the same tissue sample for P12. Scale bars: 4 mm (whole image, Left) and 50 μm (zoom, Right). (B) In P17, distinct SCLC and LCNCE components localized in separate regions of the same tissue specimen. Scale bars: 4 mm (whole image, Left) and 50 μm (zoom, Right).

**Figure S2.**
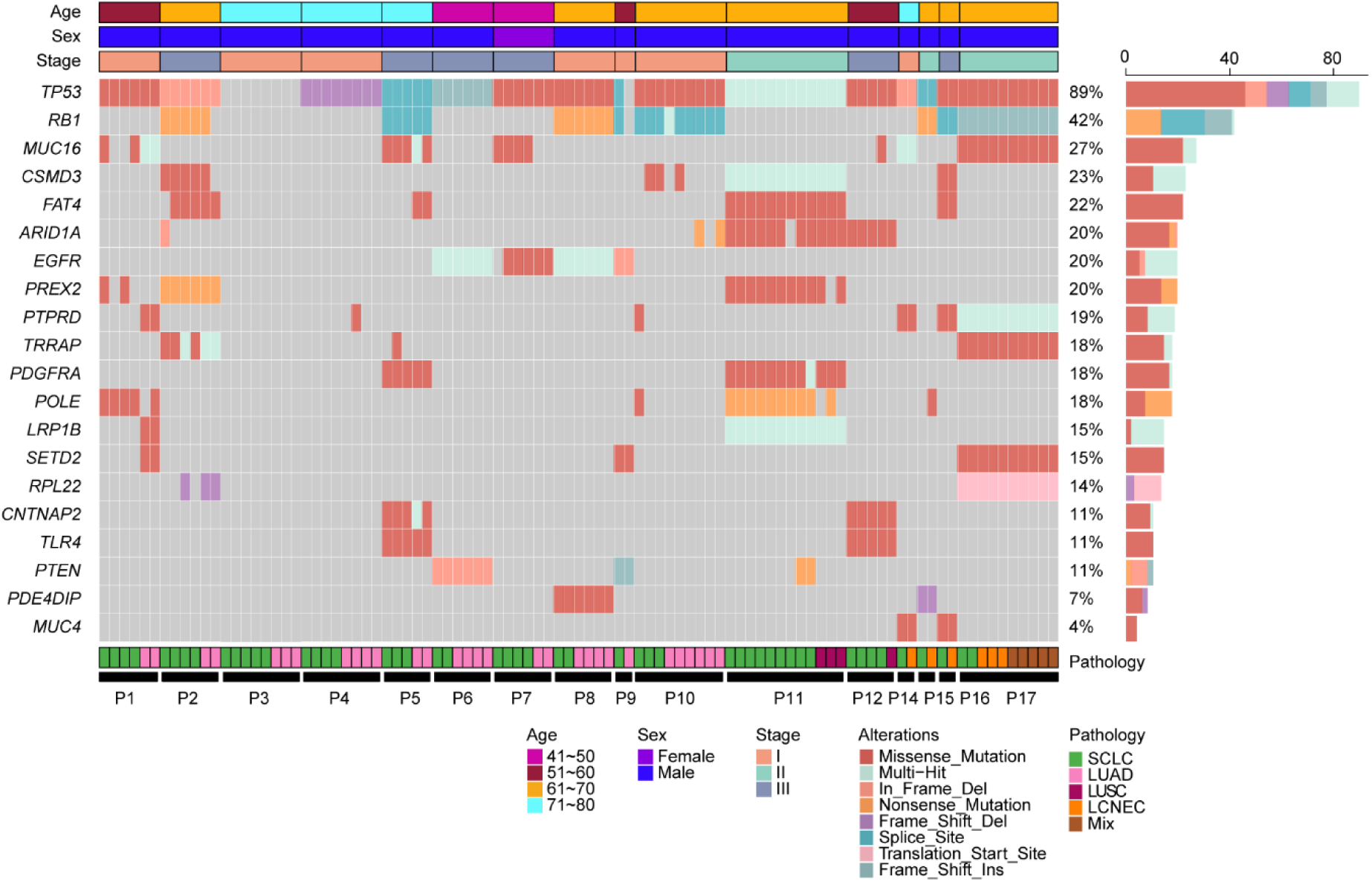
Landscape of frequent SNV and InDel on clonal mutated genes in different regions from 16 cSCLC patients. Oncoprint showing somatic alterations detected across sequenced regions from n = 16 patients with combined small cell lung cancer (cSCLC). Columns represent individual sequenced regions, grouped by patient (P1–P17 as labeled). Top annotations indicate age group (41–50, 51–60, 61–70, 71–80 years), sex (female, male), and clinical stage (I–III). The central matrix depicts alteration types for each gene and region, with colors denoting variant classes (missense mutation, nonsense mutation, splice site, translation start site, in-frame deletion, frameshift insertion/deletion) and multi-hit events (see Methods/legend for definition). Bottom annotation indicates the histological subtype of each region: small cell lung carcinoma (SCLC), lung adenocarcinoma (LUAD), lung squamous cell carcinoma (LUSC), large-cell neuroendocrine carcinoma (LCNEC), or mixed. The right bar plot shows the alteration frequency of each gene across the regions with percentages indicated.

**Figure S3.**
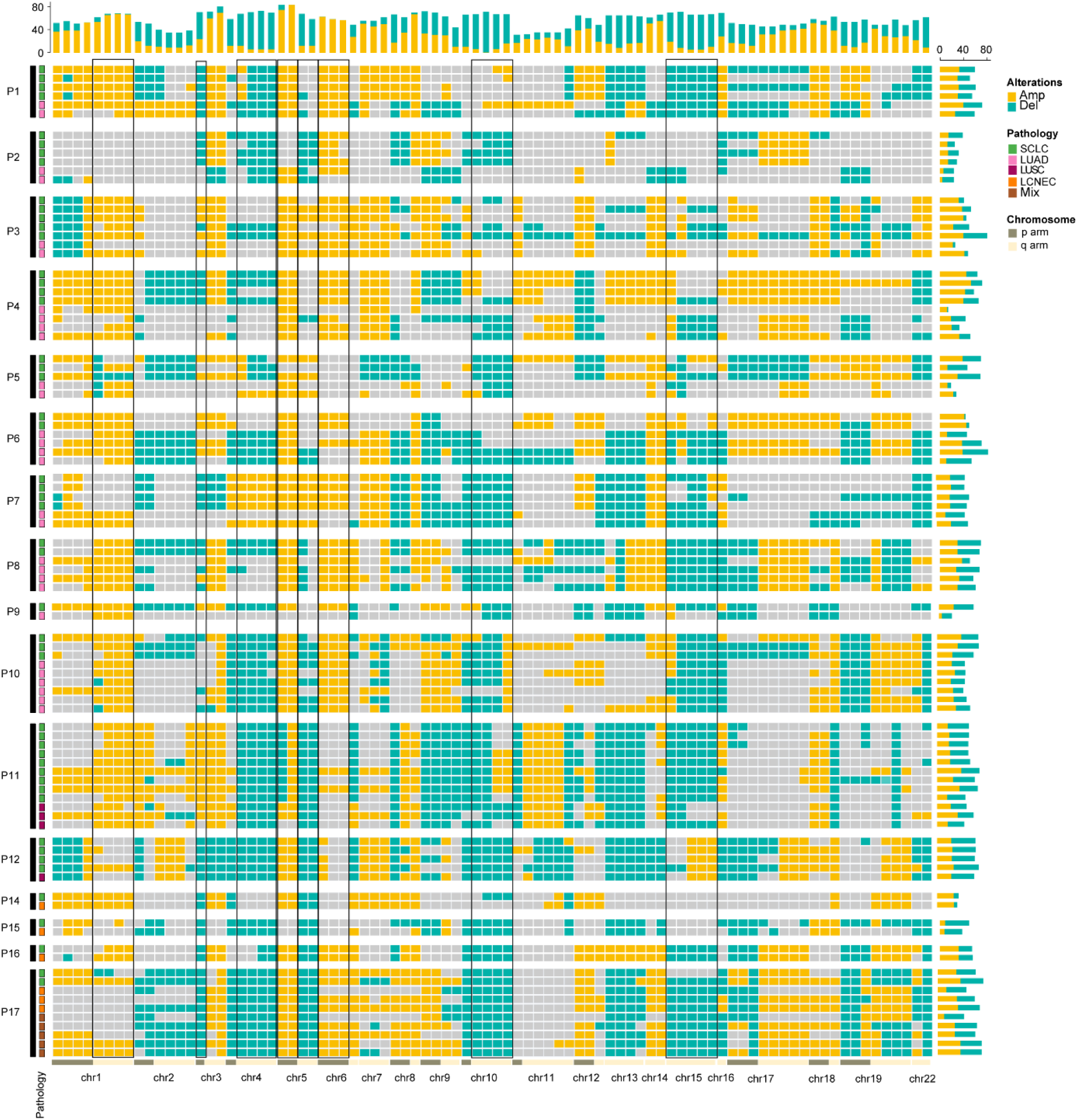
Genome-wide somatic copy-number alteration (CNA) landscape across multiregion WES samples from cSCLC patients. Heatmap showing arm-level CNA across multiple tumor regions from n = 16 cSCLC patients. Rows represent individual tumor regions (grouped by patient, P1–P17), and columns represent chromosome arms (1p–22q). CNA are color-coded as amplifications (yellow) and deletions (teal); gray indicates no call. The pathology track (left) denotes the histological subtype of each sequenced region: small cell lung cancer (SCLC), lung adenocarcinoma (LUAD), lung squamous cell carcinoma (LUSC), large-cell neuroendocrine carcinoma (LCNEC), or mixed histology (Mix). The top bar plot summarizes the frequency of CNA for each chromosome arm across all analyzed regions, with bar height indicating the percentage of regions harboring amplifications (yellow) or deletions (teal). The right bar plot shows the CNA burden for each region, expressed as the fraction of chromosome arms altered, stratified by amplifications (yellow) and deletions (teal). Black frames indicate recurrently altered chromosome arms across regions, including representative gains (e.g., 1q, 5p) and losses (e.g., 3p, 4q, 5q).

**Figure S4.**
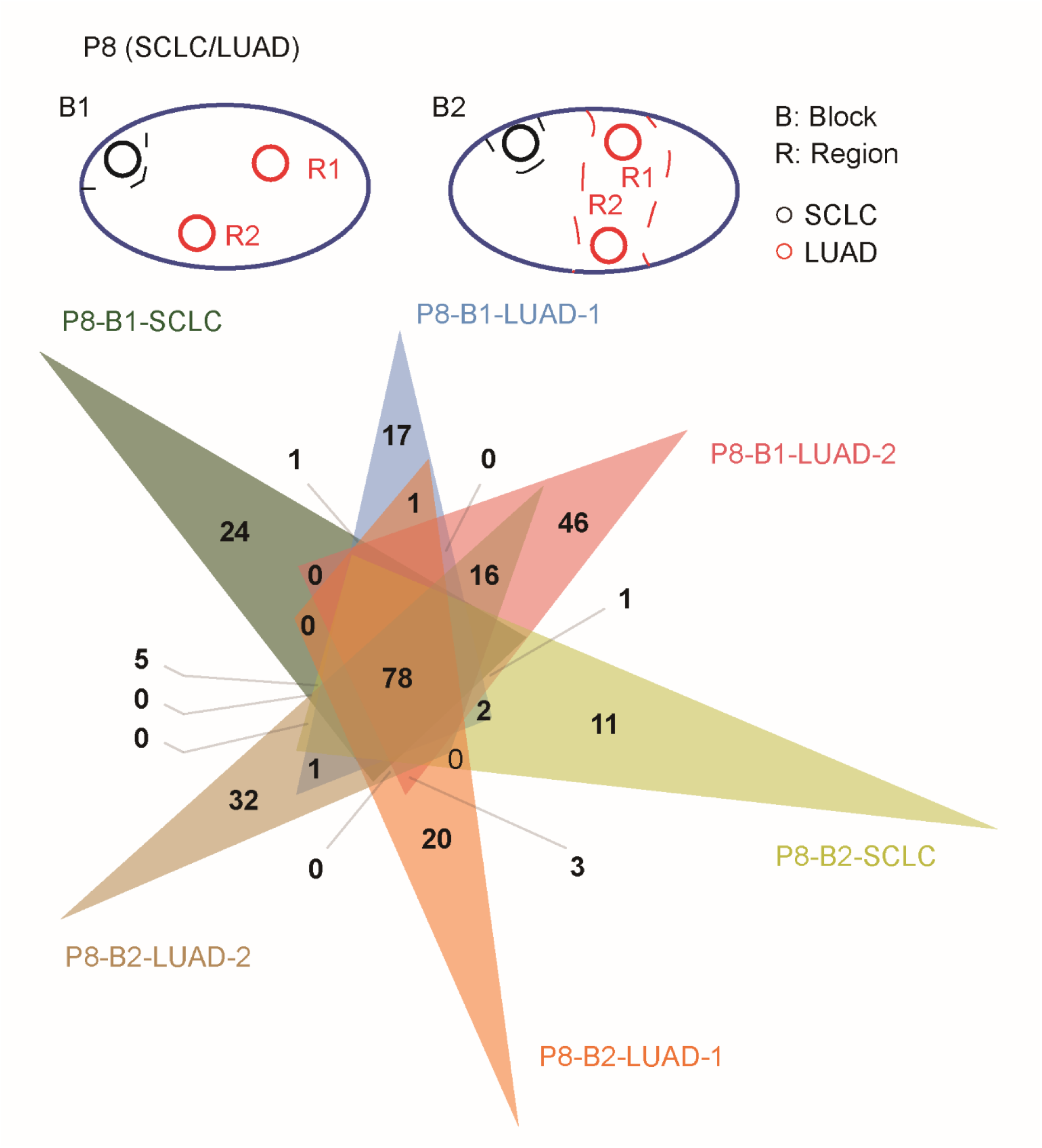
Consistency of somatic mutations across different spatial regions in P8. Schematic illustration of spatial tumor regions of SCLC or LUAD components isolated by LCM for WES in P8. Venn diagram of somatic mutations in different regions and tumor blocks, including 2 SCLC regions and 4 LUAD regions from 2 tissue blocks.

**Figure S5.**
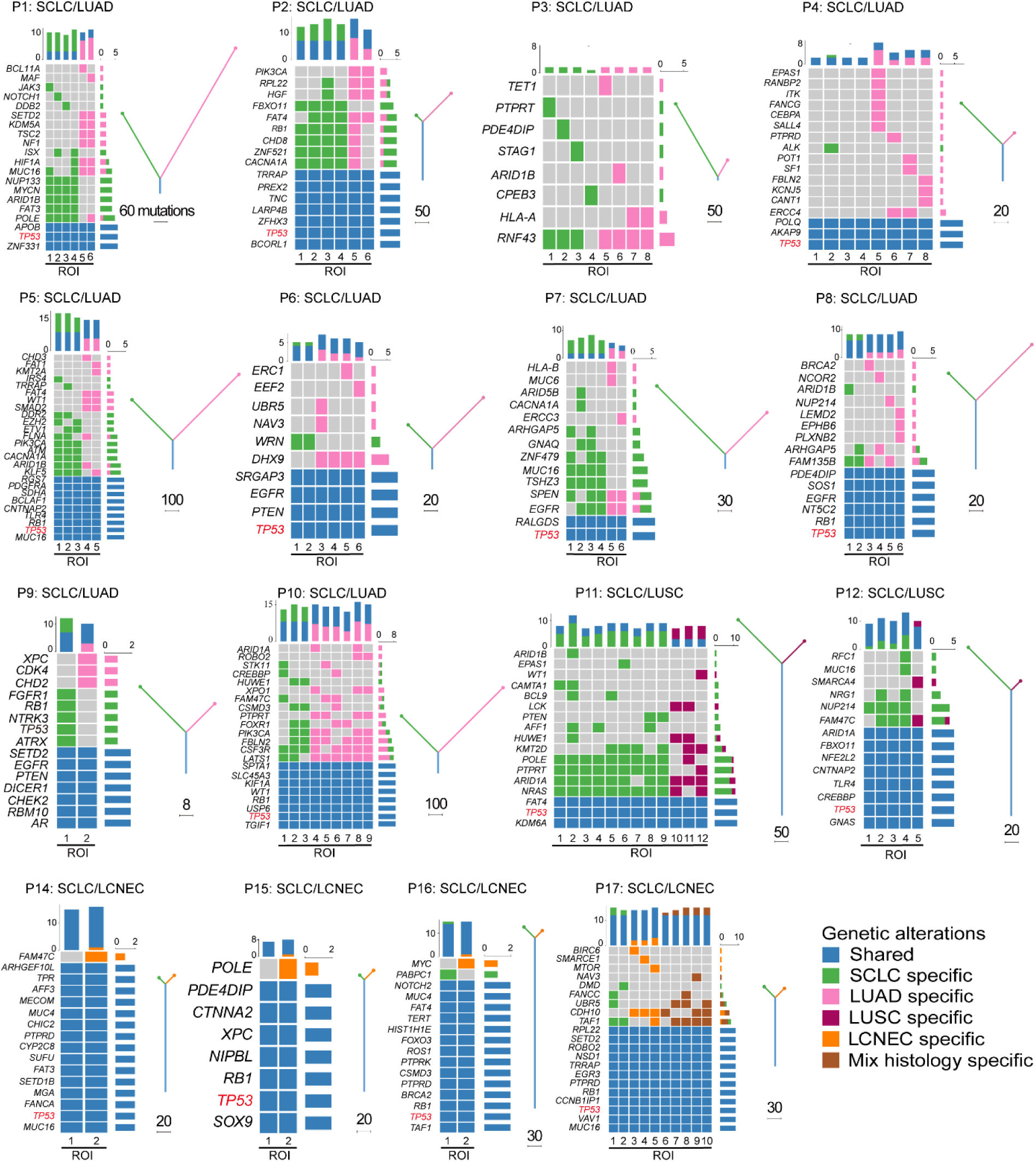
Phylogenetic trees and evolution models of 16 cSCLC patients derived from multi-region WES. For each patient panel, the central heatmap summarizes nonsynonymous somatic mutations in selected cancer genes across regions of interest (ROIs), with rows representing genes and columns representing ROIs. Mutations are color-coded by regional distribution and histology: shared/truncal (present in all sampled ROIs; blue), SCLC-specific (restricted to SCLC ROIs; green), lung adenocarcinoma (LUAD)-specific (pink), lung squamous cell carcinoma (LUSC)-specific (purple), large-cell neuroendocrine carcinoma (LCNEC)-specific (orange) and mixed-histology-specific (present only in mixed ROIs and not ubiquitous; brown). Stacked bar plots above each heatmap indicate the mutational burden per ROI (total number of nonsynonymous somatic mutations), partitioned by the same specificity categories. Phylogenetic trees shown to the right depict the inferred evolutionary relationships among ROIs; the trunk denotes early clonal mutations shared across all ROIs, whereas branches represent lineage-specific diversification associated with distinct histological components (branch colors match the corresponding pathology). Numbers labeled on trunks (e.g., “60 mutations”) indicate the count of truncal mutations. Driver genes highlighted in red (e.g., *TP53*) denote representative truncal events.

**Figure S6.**
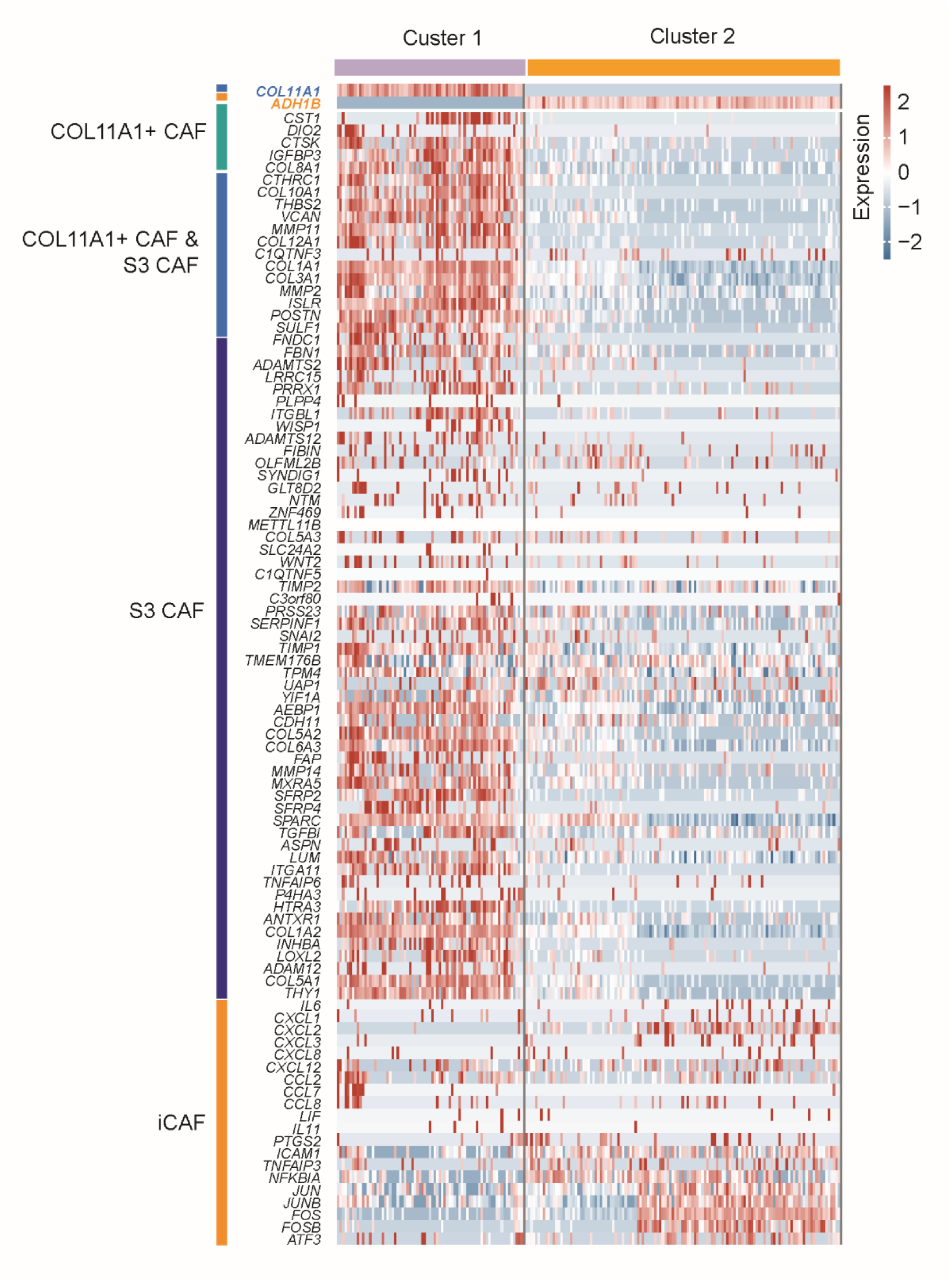
Spatial transcriptomics clusters show distinct CAF programs defined by curated COL11A1⁺ CAF and inflammatory CAF (iCAF) signature genes.^1^. Heatmap of spatial spots grouped by unsupervised clusters (Cluster 1 and Cluster 2), displaying curated marker genes for COL11A1⁺ cancer-associated fibroblasts (CAFs) and iCAFs as reported previously.**^1^**The left annotation denotes gene sets associated with COL11A1⁺ CAF (teal), S3 CAF (purple), and iCAF (orange) subtypes. Cluster 1 shows coordinated upregulation of extracellular matrix (ECM)/collagen and remodeling genes, whereas cluster 2 is enriched for inflammatory CAF genes (e.g., cytokine/chemokine and NF-κB–responsive genes). The color scale represents z-scored expression of each gene across spots.

**Figure S7.**
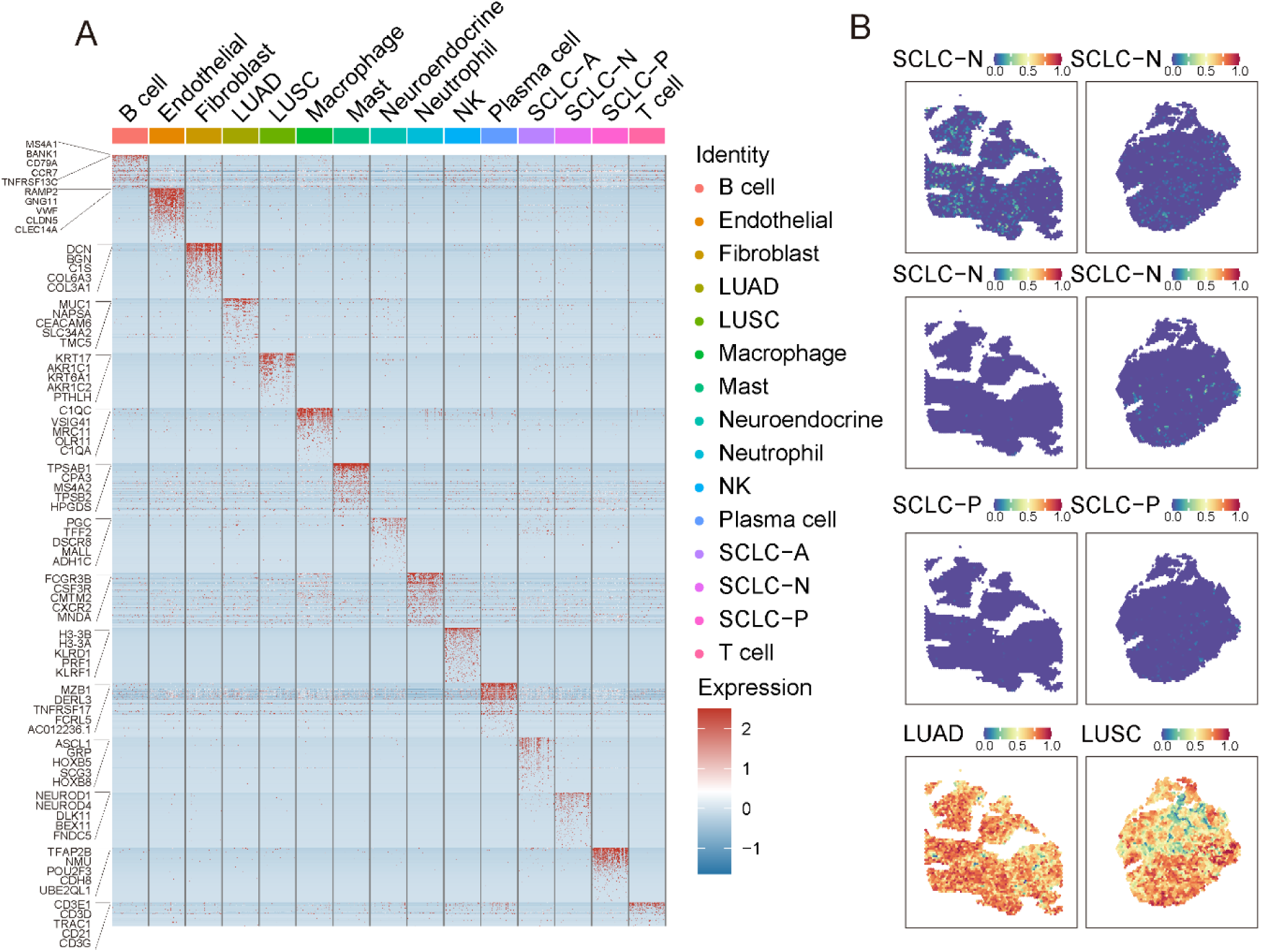
Selection and validation of cell type-specific marker genes for spatial transcriptomics deconvolution. (A) Heatmap displaying the expression profiles of the selected marker genes used to distinguish reference cell types in the ST deconvolution analysis. The reference set includes SCLC subtypes (SCLC-A, SCLC-N, SCLC-P), NSCLC subtypes (LUAD, LUSC), neuroendocrine cells, and major microenvironmental components (B cells, endothelial cells, fibroblasts, macrophages, mast cells, neutrophils, NK cells, plasma cells, and T cells). (B) Validation of the deconvolution strategy using public NSCLC spatial transcriptomics datasets.^2^ The spatial feature plots show the estimated abundance of specific tumor signatures, demonstrating that the selected markers accurately identify NSCLC components consistent with pathological evaluation, with high specificity against SCLC subtypes.

**Figure S8.**
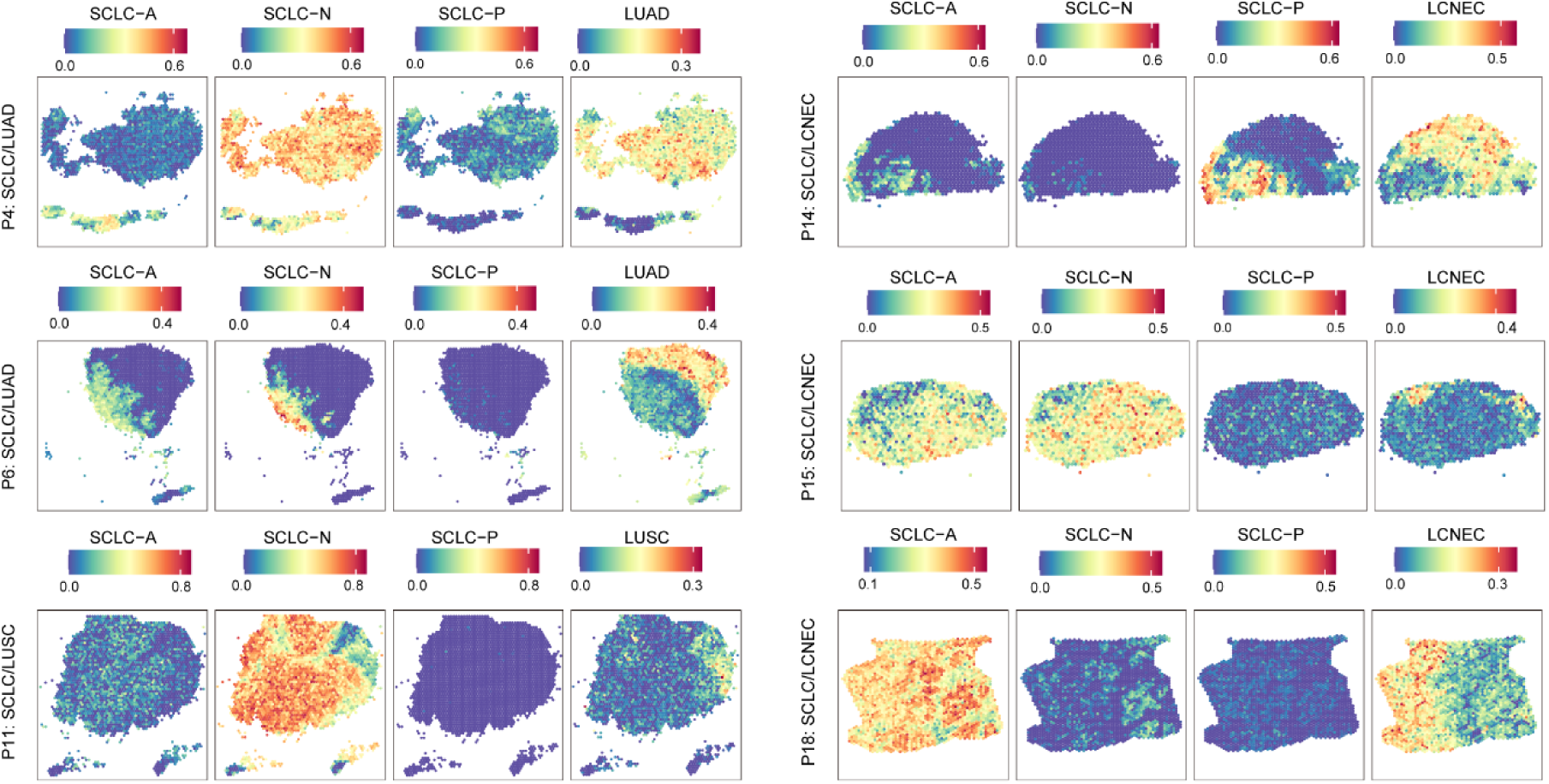
ST deconvolution inferred tumor proportions in each tumor section. SCLC-A: SCLC *ASCL1* subtype cells, SCLC-N: SCLC *NEUROD1* subtype cells, SCLC-N: SCLC *POU2F3* subtype cells, LUAD: lung adenocarcinoma, LUSC: lung squamous cell carcinoma, LCNEC: large cell neuroendocrine carcinoma

**Figure S9.**
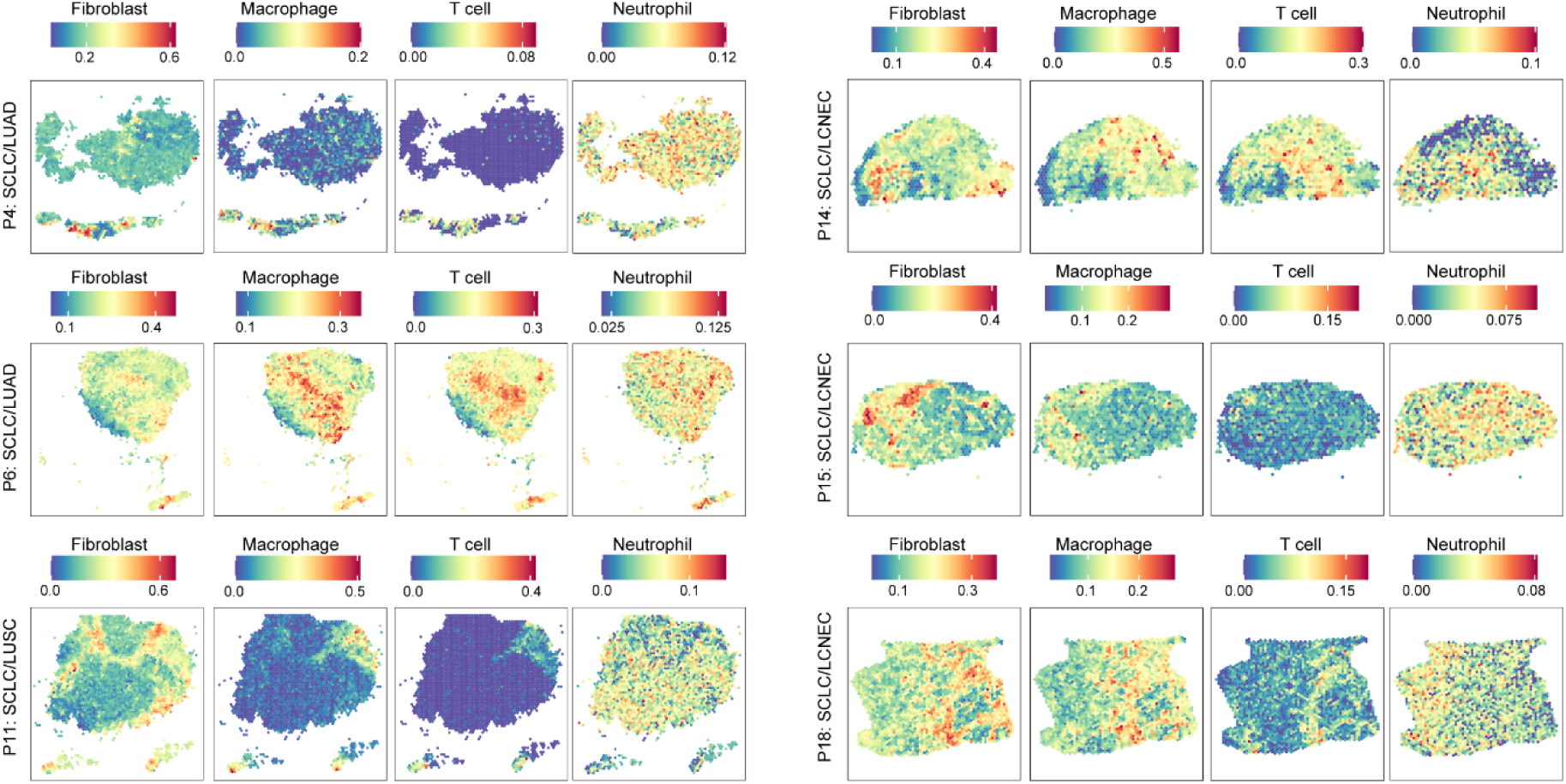
ST deconvolution inferred infiltrating cell (fibroblast, macrophage, T cell and neutrophil cell) proportions in each tumor section.

**Figure S10.**
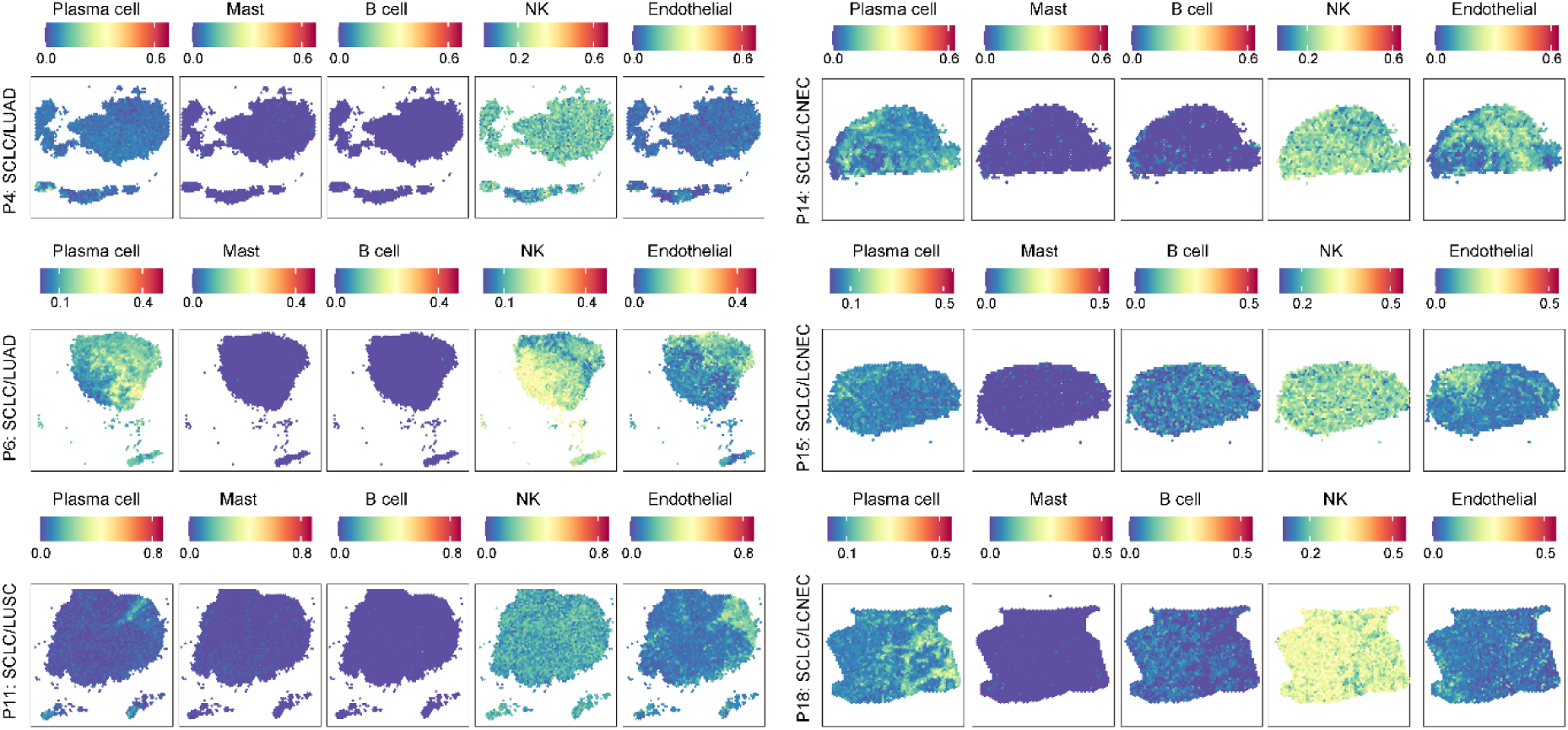
ST deconvolution inferred infiltrating cell (plasma cell, mast cell, B cell, NK cell and endothelial cell) proportions in each tumor section.

**Figure S11.**
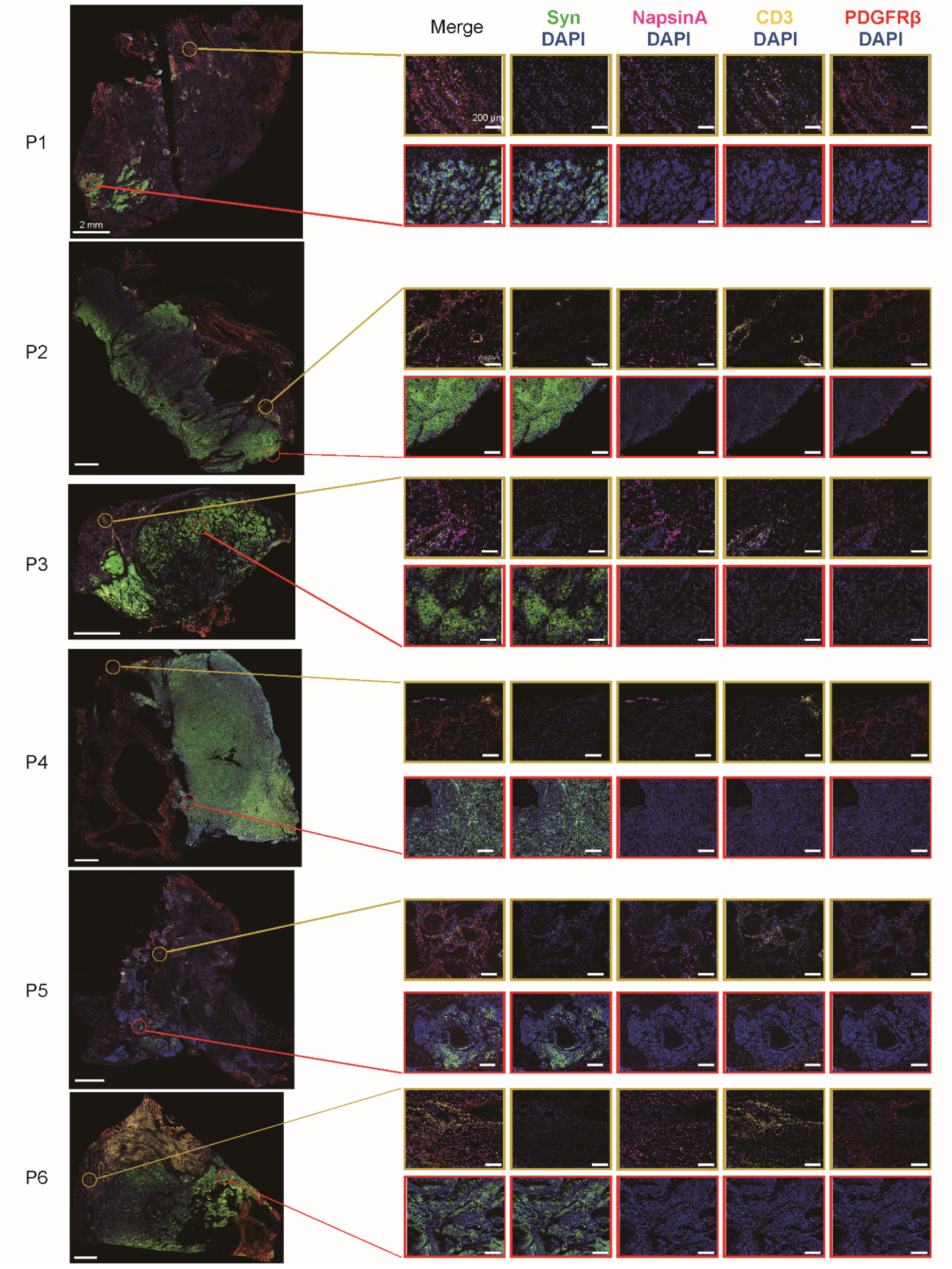
Validation of spatial TME heterogeneity *via* multiplex immunofluorescence in additional cSCLC patients (P1 to P6). Synaptophysin (Syn) for SCLC cells, Napsin A for LUAD cells, CD3 for T cells, and PDGFRβ for cancer-associated fibroblasts (CAFs). Yellow boxes indicate Napsin A^+^ LUAD domains, which are characterized by substantial infiltration of CD3^+^ T cells and PDGFRβ^+^ CAFs. Red boxes indicate synaptophysin^+^ SCLC domains, which exhibit a "cold" immune phenotype with sparse T cell and fibroblast infiltration. Scale bars: 2 mm (whole image, 0.6×magnification, Left) and 200 μm (zoom, 20× magnification, Right).

**Figure S12.**
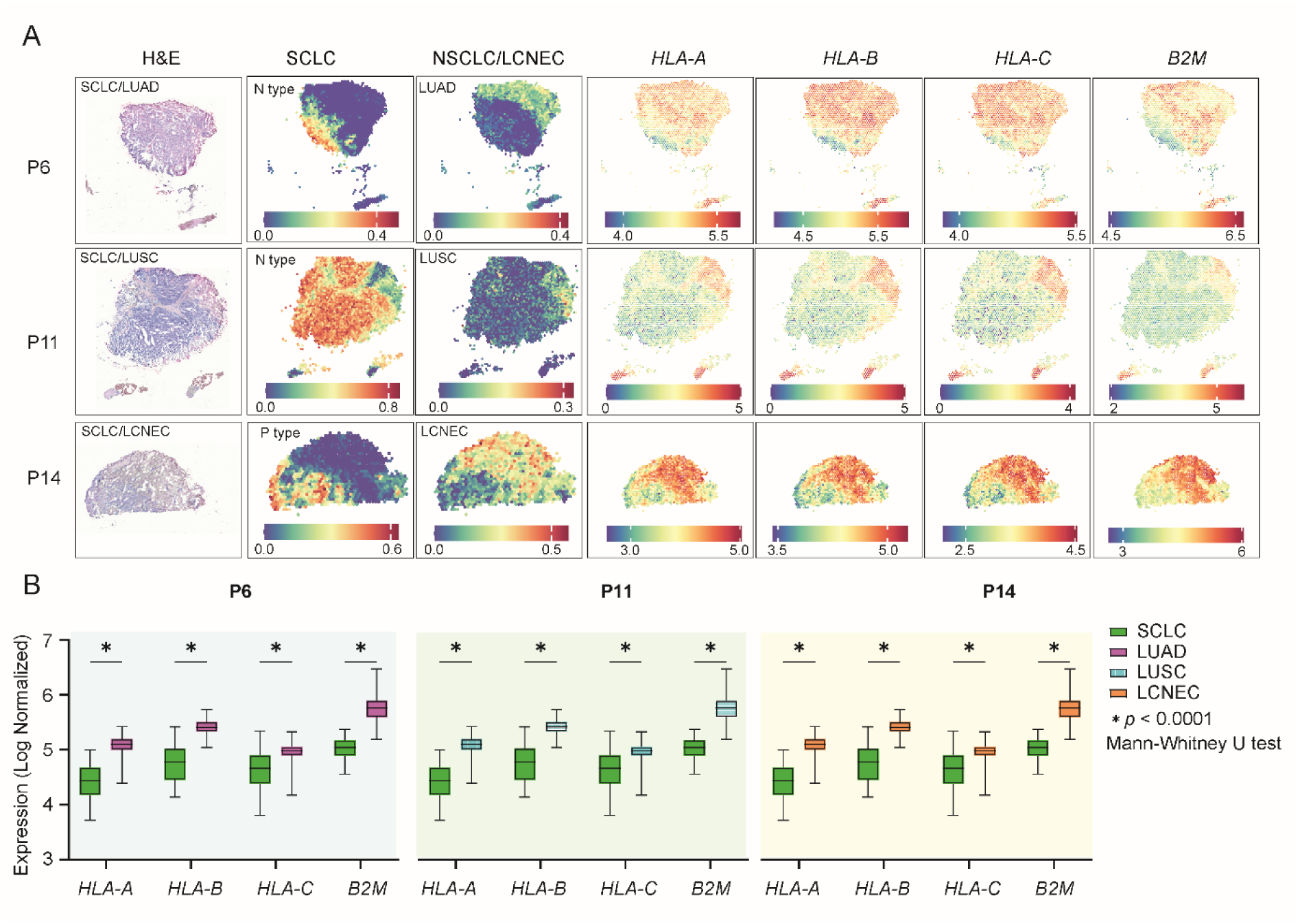
Spatial image showing *in situ* gene expression of immune infiltration related genes in the tumor section. (A) Expression of *HLA-A*, *HLA-B*, *HLA-C* and *B2M* in P6 (SCLC/LUAD), P11 (SCLC/LUSC) and P14 (SCLC/LCNEC). (B) Comparison of log-normalized expression levels of MHC Class I genes (*HLA-A*, *HLA-B*, *HLA-C*) and *B2M* between SCLC and corresponding NSCLC/LCNEC components within the same tumor sections was shown in the box plots (whisker), the central line represents the median, the box limits indicate the interquartile range (25th to 75th percentiles), and the whiskers extend to the minimum and maximum values. Statistical significance was assessed using a two-sided Mann-Whitney U test. P values were adjusted using the two-stage step-up method of Benjamini, Kriger, and Yekutieli for multiple comparison correction. (* Adjusted *p* < 0.0001).

**Figure S13.**
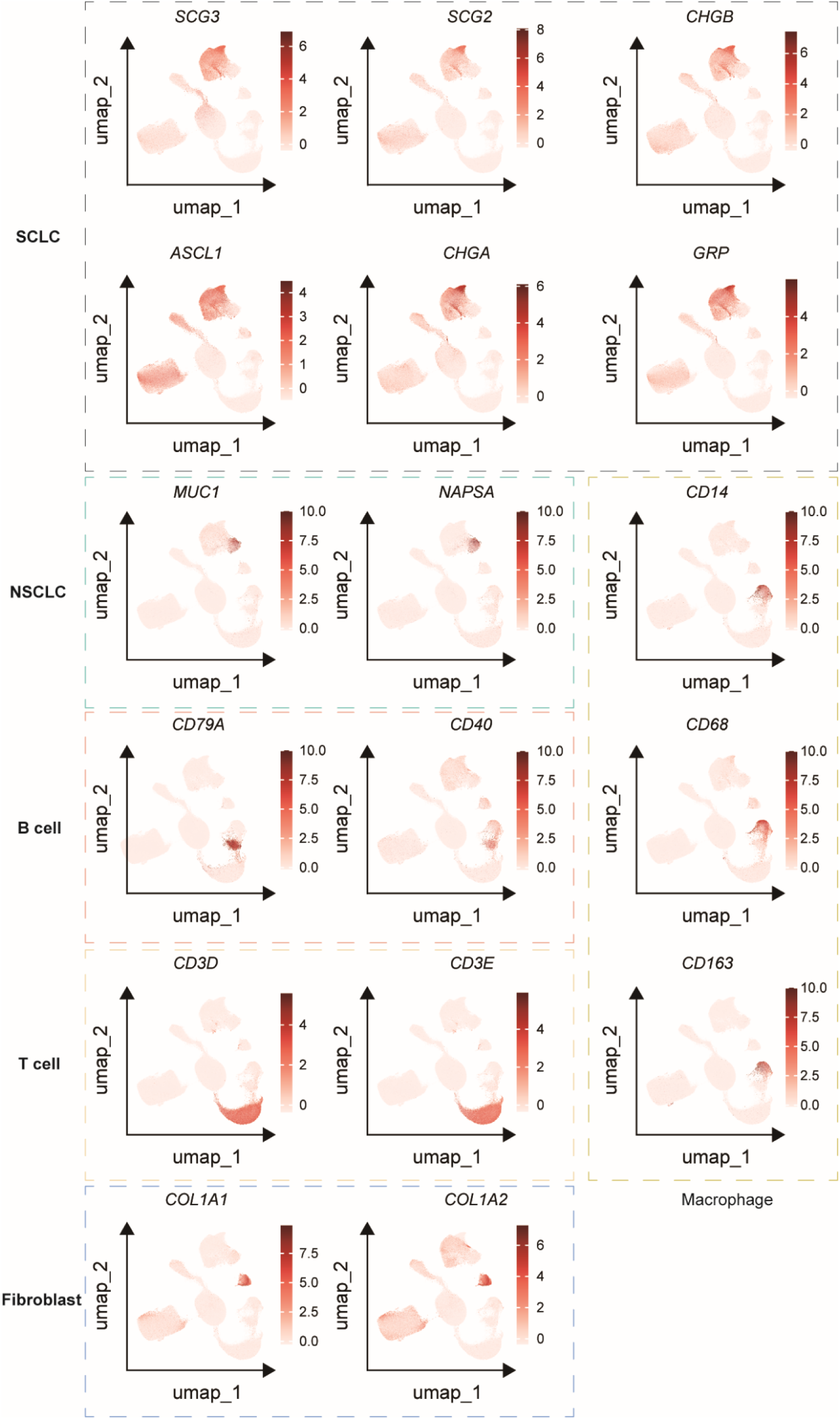
Selected marker gene expressions within the UMAP, related to Figure 7A.

**Figure S14.**
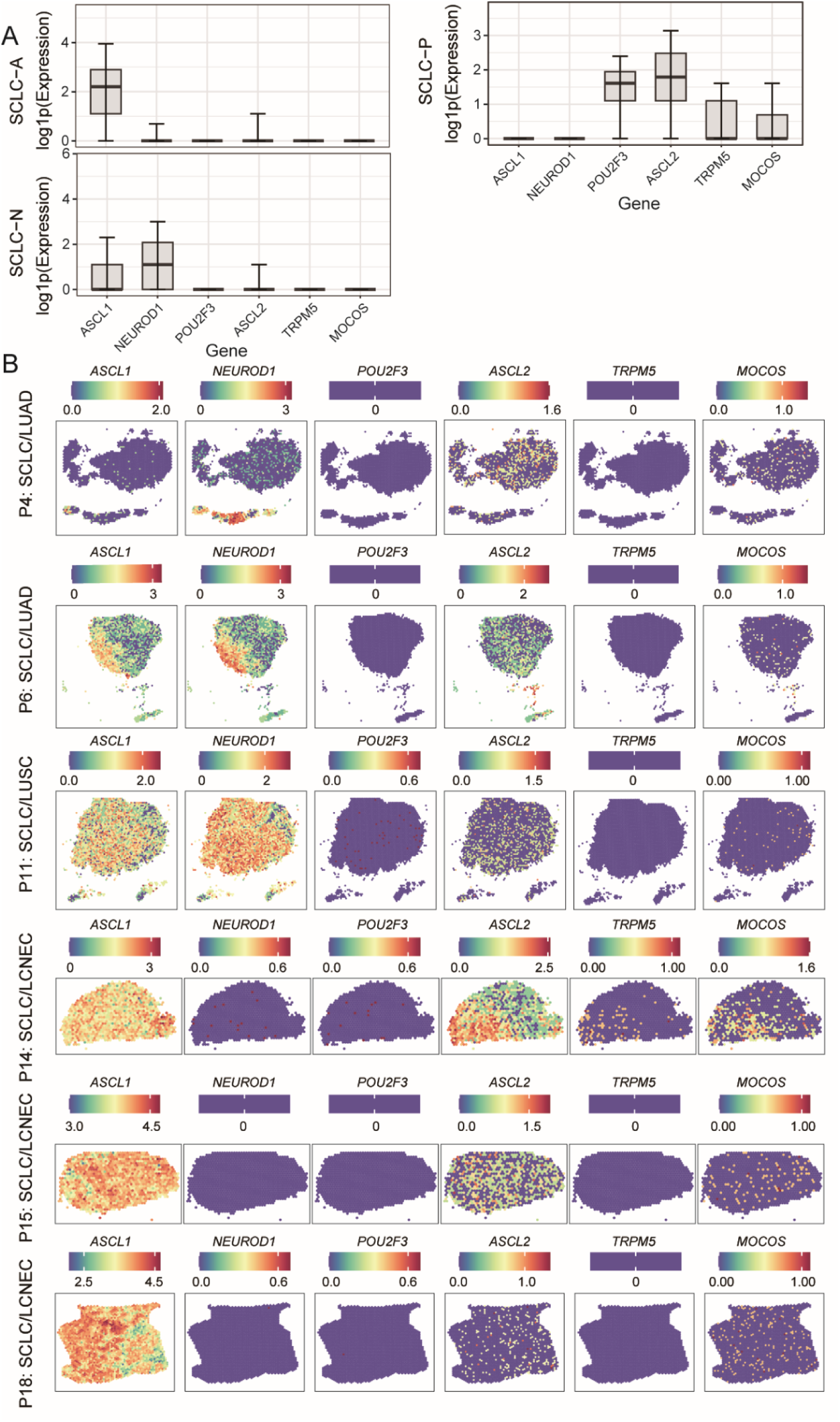
Gene expression of SCLC marker genes. (A) Boxplot showing expression of selected SCLC marker genes in SCLC *ASCL1* (SCLC-A), SCLC *NEUROD1* (SCLC-N), and SCLC *POU2F3* (SCLC-P) types, respectively. Data was shown in the box plots (whisker), the central line represents the median, the box limits indicate the interquartile range (25th to 75th percentiles), and the whiskers extend to the most extreme data point within 1.5 × IQR from the box. (B) Spatial image showing *in situ* gene expression of selected marker genes in the tumor section.

**Figure S15.**
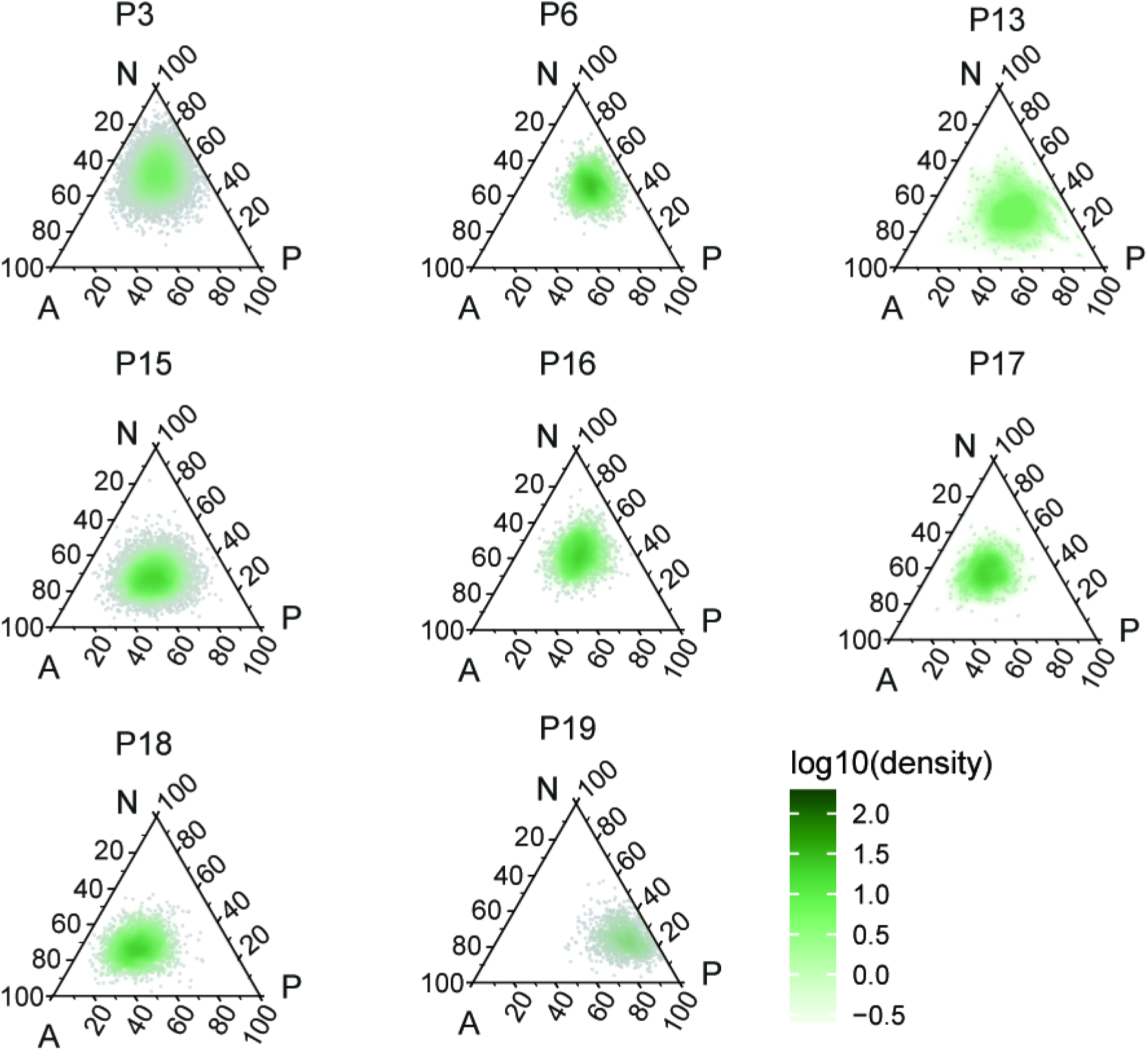
Ternary plots of individual cSCLC samples.

**Figure S16.**
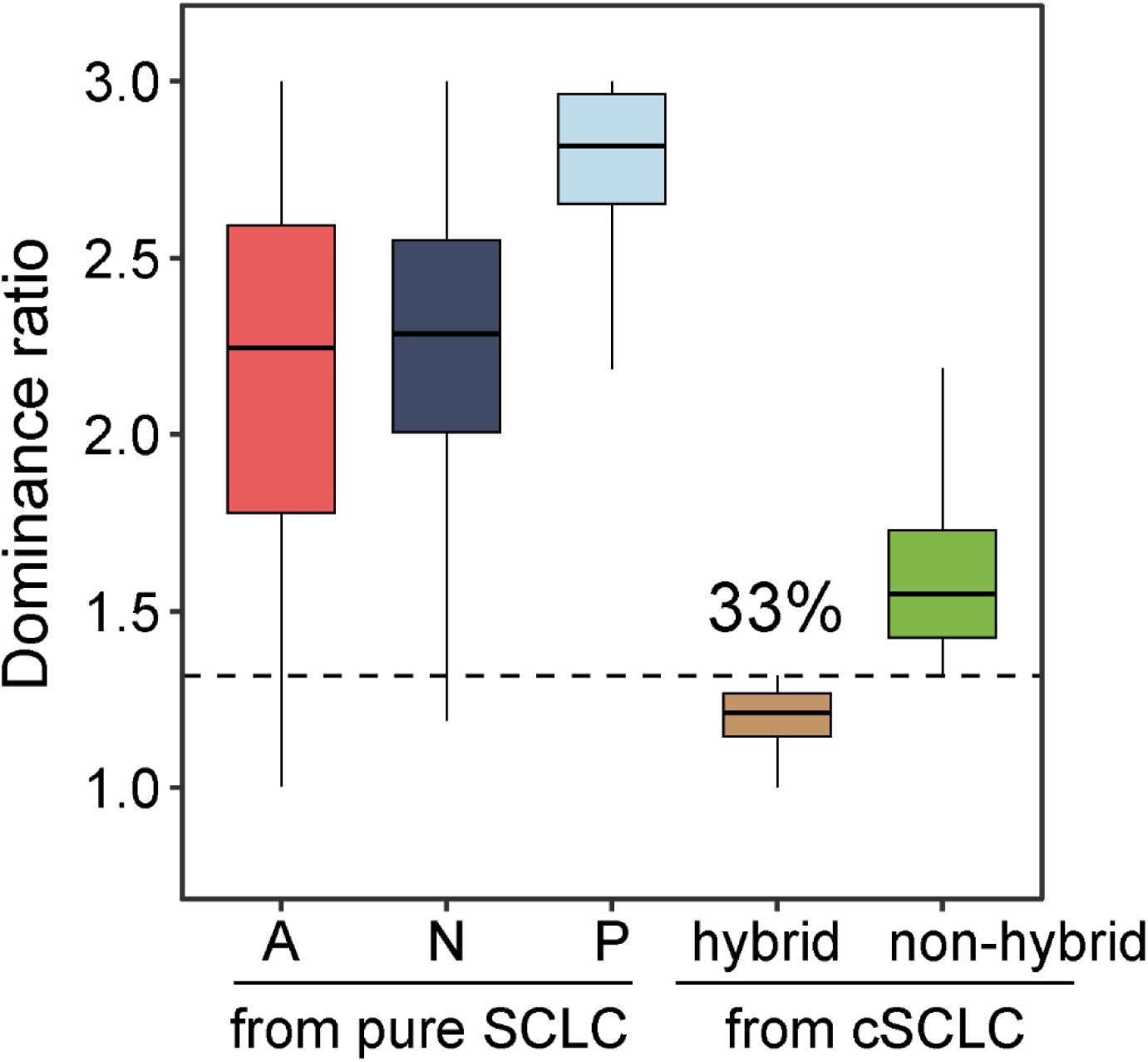
Quantification of subtype plasticity and identification of hybrid cells in cSCLC *via* dominance ratio analysis. The box plot displays the distribution of dominance ratios for reference cells from pure SCLC (subtypes A, N, and P) and SCLC-like tumor cells from cSCLC samples. Data was shown in the box plots (whisker), the central line represents the median, the box limits indicate the interquartile range (25th to 75th percentiles), and the whiskers extend to the most extreme data point within 1.5 × IQR from the box.

**Figure S17.**
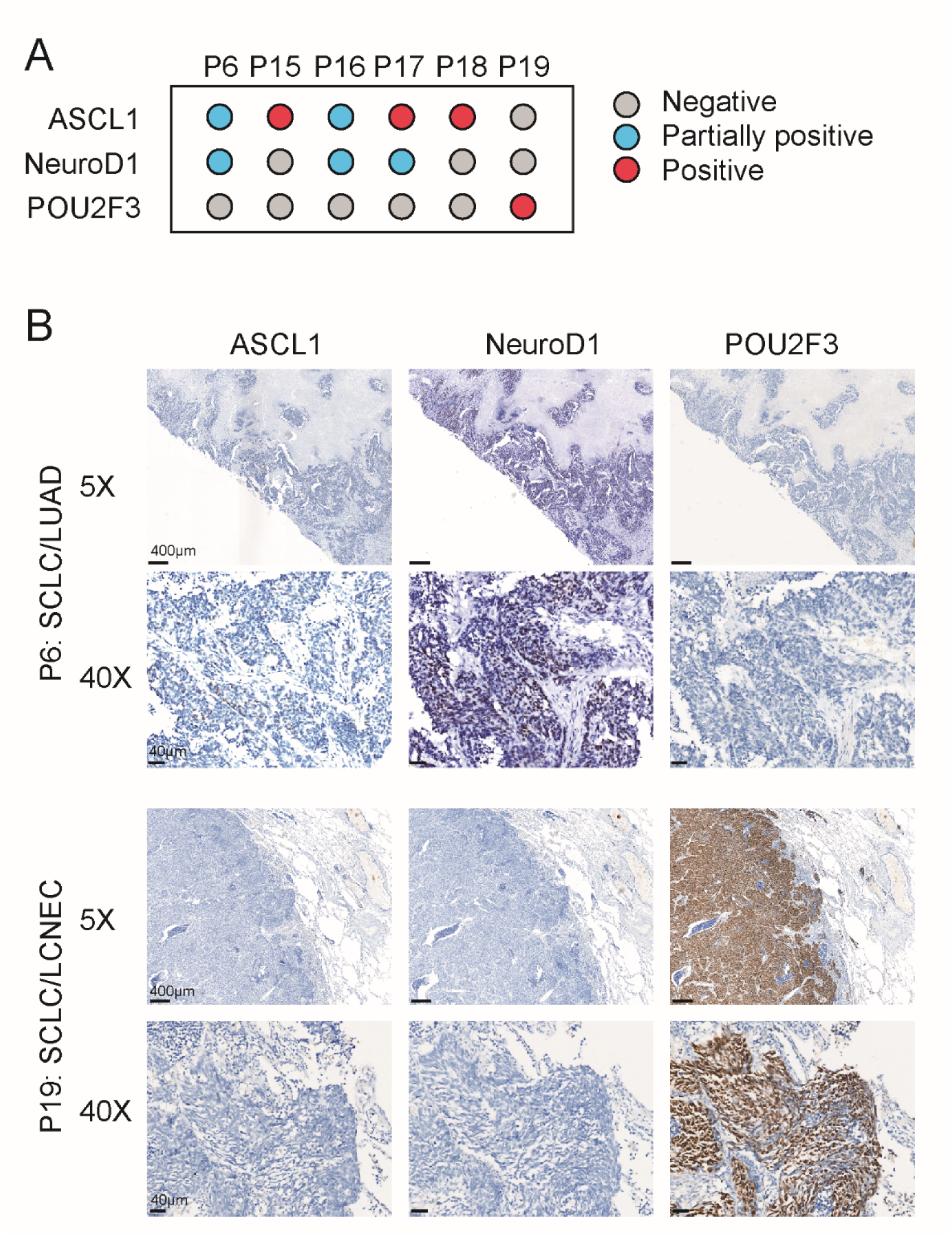
Immunohistochemical validation of SCLC lineage transcription factors in cSCLC patients. (A) Summary of immunohistochemical (IHC) staining results for three key SCLC lineage transcription factors-ASCL1, NeuroD1, and POU2F3-across six selected cSCLC patients (P6, P15-P19). (B) Representative IHC micrographs from patient P6 (SCLC/LUAD subtype) and patient P19 (SCLC/LCNEC subtype). The panels display expression patterns of ASCL1, NeuroD1, and POU2F3. P6 exhibits partial positivity for ASCL1 and NeuroD1, consistent with the SCLC-A/N hybrid features, while P19 shows exclusive and strong positivity for POU2F3, confirming the SCLC-P subtype. Scale bars: 400 μm (5X magnification) and 40 μm (40X magnification).

### Supplementary Tables

**Table S1.** Demographics and clinical characteristics of cSCLC patients in the cSCLC cohort, as well as the summary of multi-omic characterization of cSCLCs.

| Patient | Sex | Age | Histopathology | No. of tissue blocks | Stage | Smoking | WES | ST | snRNAseq |
| --- | --- | --- | --- | --- | --- | --- | --- | --- | --- |
| P1 | male | 59 | SCLC/LUAD | 1 | I | Nonsmoker | Yes | / |  |
| P2 | male | 66 | SCLC/ LUAD | 1 | III | Unknown | Yes | / |  |
| P3 | male | 68 | SCLC/ LUAD | 2 | I | Smoker | Yes | / | Yes |
| P4 | male | 73 | SCLC/ LUAD | 2 | I | Smoker | Yes | Yes | Yes |
| P5 | male | 72 | SCLC/ LUAD | 2 | III | Nonsmoker | Yes | / |  |
| P6 | male | 42 | SCLC/ LUAD | 2 | III | Smoker | Yes | Yes | Yes |
| P7 | female | 47 | SCLC/ LUAD | 2 | III | Nonsmoker | Yes | / |  |
| P8 | male | 65 | SCLC/ LUAD | 2 | I | Nonsmoker | Yes | / |  |
| P9 | male | 55 | SCLC/ LUAD | 2 | I | Smoker | Yes | / |  |
| P10 | male | 66 | SCLC/ LUAD | 3 | I | Smoker | Yes | / |  |
| P11 | male | 65 | SCLC/LUSC | 3 | II | Nonsmoker | Yes | Yes | Yes |
| P12 | male | 57 | SCLC/LUSC | 2 | III | Smoker | Yes | / | Yes |
| P13 | male | 73 | SCLC/LUSC | 1 | II | Nonsmoker | / | / | Yes |
| P14 | male | 78 | SCLC/LCNEC | 1 | I | Unknown | Yes | Yes | Yes |
| P15 | male | 69 | SCLC/LCNEC | 1 | II | Unknown | Yes | Yes | Yes |
| P16 | male | 61 | SCLC/LCNEC | 1 | III | Unknown | Yes | / | Yes |
| P17 | male | 64 | SCLC/LCNEC | 2 | II | Unknown | Yes | / | Yes |
| P18 | male | 65 | SCLC/LCNEC | 1 | I | Unknown | / | Yes | Yes |
| P19 | male | 68 | SCLC/LCNEC | 1 | III | Unknown | / | / | Yes |

**Table S2.** Quality control of multi-region WES and the numbers of somatic mutations detected in the cSCCL cohort (see separate file of spreadsheets).

**Table S3.** Information of somatic mutations detected in the cSCCL cohort (see separate file of spreadsheets).

**Table S4.** Information of *TP53* gene mutations across tumor components (see separate file of spreadsheets).

**Table S5.**
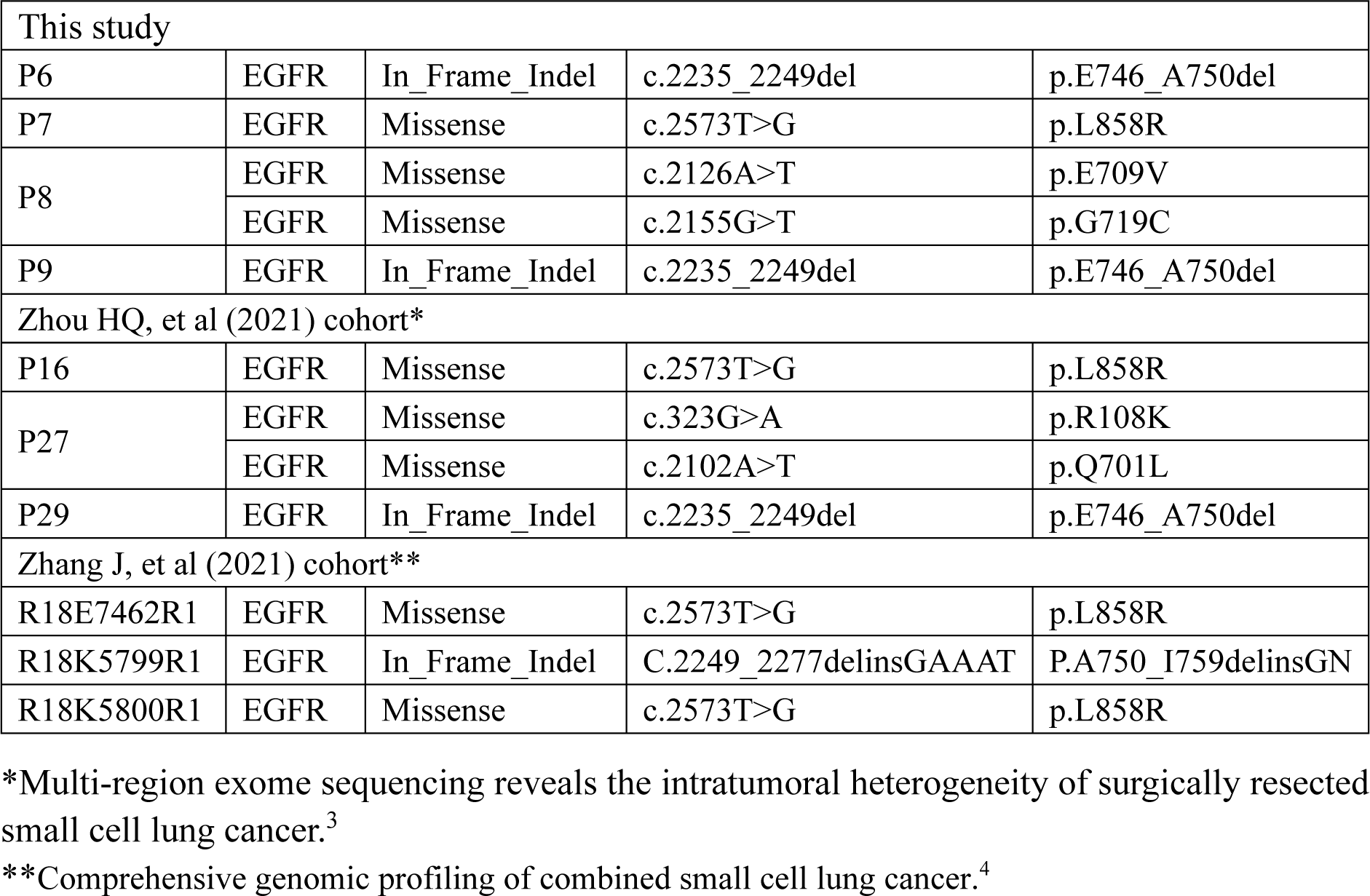
*EGFR* mutations of cSCLC patients in this study and public datasets.

| This study |  |  |  |  |
| --- | --- | --- | --- | --- |
| P6 | EGFR | In_Frame_Indel | c.2235_2249del | p.E746_A750del |
| P7 | EGFR | Missense | c.2573T>G | p.L858R |
| P8 | EGFR | Missense | c.2126A>T | p.E709V |
|  | EGFR | Missense | c.2155G>T | p.G719C |
| P9 | EGFR | In_Frame_Indel | c.2235_2249del | p.E746_A750del |
| Zhou HQ, et al (2021) cohort* |  |  |  |  |
| P16 | EGFR | Missense | c.2573T>G | p.L858R |
| P27 | EGFR | Missense | c.323G>A | p.R108K |
|  | EGFR | Missense | c.2102A>T | p.Q701L |
| P29 | EGFR | In_Frame_Indel | c.2235_2249del | p.E746_A750del |
| Zhang J, et al (2021) cohort** |  |  |  |  |
| R18E7462R1 | EGFR | Missense | c.2573T>G | p.L858R |
| R18K5799R1 | EGFR | In_Frame_Indel | C.2249_2277delinsGAAAT | P.A750_I759delinsGN |
| R18K5800R1 | EGFR | Missense | c.2573T>G | p.L858R |

\*Multi-region exome sequencing reveals the intratumoral heterogeneity of surgically resected small cell lung cancer.<sup>3</sup>
\*\*Comprehensive genomic profiling of combined small cell lung cancer.<sup>4</sup>

**Table S6.** Information of gene mutations for cSCLC Detector in public cSCLC cohorts (see separate file of spreadsheets).

**Table S7.** Information of gene mutations and samples in public cSCLC cohorts (see separate file of spreadsheets).

